# Genetically Defined Organoid Models Reveal Mechanisms Driving Squamous Cell Neoplastic Evolution and Identify Potential Therapeutic Vulnerabilities

**DOI:** 10.1101/2025.01.18.631624

**Authors:** Hua Zhao, Young Min Park, Yueyuan Zheng, Qiong Mao, Casey Collet, Boyan Hu, Tianming Zhou, Luda Lin, Stephanie Wong, Yuhao Pan, Anette Vistoro Monreal, Uttam K. Sinha, Parish Sedghizadeh, Alice Soragni, De-Chen Lin

## Abstract

Upper aerodigestive squamous cell carcinoma (UASCC) is an aggressive and lethal neoplasm, with its early neoplastic transformation mechanisms remaining poorly understood. Here, we characterize over 25 genetically-defined organoid models derived from murine and human oral/esophageal tissues harboring key driver mutations. Double knockout of *TP53* and *CDKN2A* induced morphological dysplasia, hyperproliferation, loss of squamous differentiation, and tumorigenicity, which were further exacerbated by additional driver mutations (e.g., *PIK3CA*, *NOTCH1*, *KMT2C*). Single-cell analysis revealed an expansion of quiescent basal cells and proliferative squamous cells, alongside a loss of differentiated squamous cells during malignant transformation. A distinct senescence program, regulated by ANXA1, was markedly diminished during early neoplastic evolution. Mechanistically, the ANXA1-SMAD3-p27^KIP1^ pathway was identified as a critical regulator of this senescence program, acting to suppress neoplastic features in organoid models. Lastly, our high-throughput, single-organoid-resolution drug screens unexpectedly revealed *PIK3CA*-driven organoids exhibited sensitivity to Mitomycin C and Onalespib. This study provides novel mechanistic insights into early neoplastic evolution and underscores the value of genetically-defined organoid models for investigating cancer biology and identifying targeted therapies.

## INTRODUCTION

Upper aerodigestive squamous cell carcinoma (UASCC) is among the most lethal malignancies, causing over 1.5 million new cases and 900,000 deaths annually (1). SCCs of the head and neck (HNSCC) and esophagus (ESCC), mostly arising from the mucosal epithelium in the oral cavity, oropharynx, and esophagus, constitute over 90% of UASCC. Current treatment of UASCC is highly challenging, often involving a combination of surgery, radiation, and chemotherapy. Given the critical functions of the mouth and esophagus, this multimodal approach often results in severe side effects and long-lasting disability, significantly affecting the quality of life – often rivaling the burden of the cancer itself. High rates of resistance to conventional therapies, coupled with substantial psychological distress, highlight the urgent, unmet need for a deeper understanding of UASCC pathophysiology to improve therapeutic outcomes and patient care.

The malignant transformation of UASCC follows a multistep tumorigenesis process, beginning with benign squamous epithelium transitioning to dysplasia, progressing to carcinoma in situ, and eventually advancing to invasive SCC (2,3). Despite this well-defined paradigm, our understanding of the molecular mechanisms underlying this stepwise neoplastic evolution remains limited, significantly hindering the development of effective strategies for prevention and early intervention.

A major obstacle to advancing this research has been the lack of viable and validated models that accurately recapitulate this dynamic, pathological transition. Efforts to address these challenges include carcinogen-treated and genetically engineered mouse models (4,5). However, these approaches are time-consuming, costly, and pose additional challenges, such as anatomical differences between humans and mice, as well as the limited material and availability of murine precursor lesions for in-depth biological investigation (6,7).

The three-dimensional (3D) organoid culture system provides a robust and valid alternative for studying premalignancy and early neoplastic evolution. This system recapitulates and maintains the genetic, biological and phenotypic characteristics of corresponding tissues of origin (8,9). Moreover, organoid culturing has made it feasible and efficient to genetically manipulate the cells-of-origin directly from human samples. We have previously established human 3D organoid model systems for various tissues, such as HNSCC tumor specimens (10), Barrett’s esophagus (11), gastroesophageal junction (12), among others (9,13). Notably, we and others have shown that CRISPR/Cas9-based genome editing of normal organoids successfully models early neoplastic transformations in various human tissues (8,12,14–16).

In the present study, we established both normal and genetically-engineered murine and human organoids to recapitulate the early squamous malignant transformation process induced by key cancer drivers (*TP53*, *CDKN2A*, *PIK3CA*, *NOTCH*, *KMT2C* and *CDKN1B*). Single cell genomic analysis unexpectedly identified a senescence-like gene program as the most suppressed program during this pathological transition. We furthermore explored the potential of genetically-engineered and patient-derived tumor organoids for drug screening and validation of novel effective drugs.

## RESULTS

### A Cross-Species, Genetically-Engineered Organoid System to Model Early Neoplastic Evolution of UASCC

*TP53* mutations (86-93% patients) and *CDKN2A* mutations/deletions (73-78% patients) are the most common drivers in UASCC (17,18) (**Fig. 1A**). Importantly, loss of these two drivers occurs early during the neoplastic evolution. For example, the most common genomically inactivated genes in oral leukoplakia samples (precursor lesions to HNSCC) are *TP53* and *CDKN2A* (19,20). Thus, to model early neoplastic transformation of UASCC, we generated normal organoid lines from oral/esophageal epithelial cells, followed by the knockout of *TP53*/*CDKN2A* using the CRISPR/Cas9 genome editing (**Fig. 1B**).

**Figure 1.**
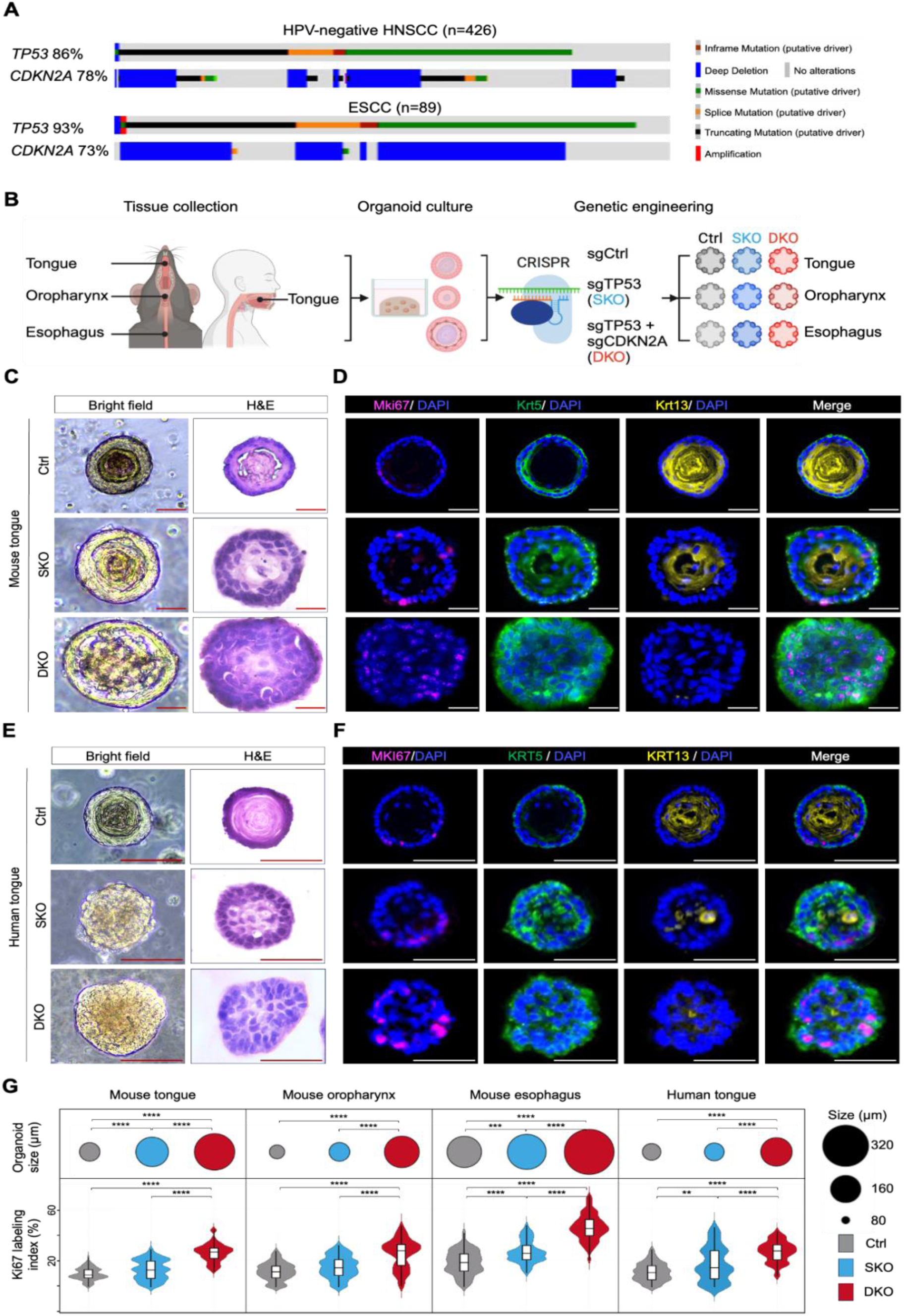
Cross-species, genetically-engineered organoid modeling early neoplastic evolution of UASCC. **(A)** Genomic aberrations of *TP53* and *CDKN2A* in HNSCC and ESCC tumors from the TCGA cohort, plotted using the Cbioportal(22). **(B)** Schematic of organoid generation and genetic engineering using CRISPR-Cas9 to create *TP53* single-knockout (SKO), *TP53* and *CDKN2A* double-knockout (DKO), and control (Ctrl) lines. This plot was generated via BioRender. **(C)** Representative bright-field, hematoxylin-and-eosin (H&E), and **(D)** immunofluorescent (IF) images showing the proliferation marker Mki67 (magenta), basal cell marker Krt5 (green), and squamous differentiation marker Krt13 (yellow) in 3-week-old mouse tongue Ctrl, SKO, and DKO organoids. (**E)** Representative bright-field, H&E, and **(F)** IF images showing staining of MKI67 (magenta), KRT5 (green), and KRT13 (yellow) in 3-week-old human tongue Ctrl, SKO, and DKO organoids. **(G)** Quantification of organoid size and Ki67 labeling index of 3-week-old genetically engineered mouse and human organoid lines (*n* = 50 per group). Scale bar, 100μm; \*\**P* < 0.01; \*\*\**P* < 0.001; \*\*\*\**P* < 0.0001.

We first refined conditions for establishing mouse normal and mutant organoid models. Freshly collected murine tongue, oropharynx, and esophageal epithelial layers were digested into a single-cell suspension and seeded in basement membrane extracts (BME) and organoid-conditioned medium. After seven days, individual organoids were harvested under microscopy, separated from debris, dissociated into single cells and reseeded (**Fig. S1A**). Three weeks post-passage, murine tongue, oropharynx, and esophageal organoids formed dense 3D structures with average diameters of 102 μm, 91 μm, and 174 μm, respectively. Immunostaining analyses showed that murine organoid recapitulated the stratified squamous epithelium histology, composed of peripheral Krt5^+^ basal cells and Krt13^+^ differentiated squamous cells, as well as full-differentiated keratin cores (**Fig. S1B-C**). These murine organoids could be expanded and propagated continuously for over 21 months ex vivo with biweekly passaging.

We next electroporated either scramble control, *Trp53*-targeting, or *Trp53*/*Cdkn2a*-dual-targeting Cas9:gRNA ribonucleoprotein (RNP) complexes into these murine organoids to generate control (Ctrl), *Trp53* single-knockout (SKO), and *Trp53*/*Cdkn2a* dual-knockout (DKO) organoid lines. High electroporation efficiency (44%-61%) was confirmed under fluorescence microscopy (**Fig. S1D**). A two-week nutlin-3a functional selection (12,21) yielded the outgrowth and expansion of SKO and DKO organoids. The presence of frameshift mutations in *Trp53* and *Cdkn2a* at the expected locations was further confirmed by TOPO cloning and Sanger sequencing (**Fig. S1E**).

To characterize the phenotypic changes upon *Trp53*-single loss or *Trp53*/*Cdkn2a*-dual loss, we initiated mouse organoid formation from a single-cell suspension and analyzed the growth properties and histological features. Compared with Ctrl mouse tongue organoids, SKO organoids showed a consistent increase in size and a higher metabolically activity till day 7 (**Fig. S2A**). By day 21, SKO organoids exhibited substantially enlarged, atypical nuclei (**Fig. 1C-D**). Moreover, SKO organoids displayed a conspicuous expansion of Krt5^+^ basal cell population, contraction of Krt13^+^ differentiated squamous cell compartment, and loss of well-differentiated keratin core (**Fig. 1D**), highlighting dysplastic features. Importantly, these phenotypic abnormalities and neoplastic characteristics were further exacerbated in the DKO organoids (**Fig. 1C-D, Fig. S2A-C**). In mouse oropharynx organoids, SKO lines exhibited atypical cell nuclei and higher nuclear-cytoplasmic ratios, while there were slight but not significant increases in terms of organoid size, cell viability, or Mki67 labeling index (**Fig. 1G, Fig. S3**). In contrast, DKO oropharynx organoids displayed strong dysplastic morphology, including hyperplastic growth and reduced suprabasal and basal differentiation features compared to Ctrl organoids (**Fig. 1G, Fig. S3**). In the context of esophageal organoids, SKO notably induced architectural complexity besides hyperproliferation, hyperplastic Krt5^+^ basal cells, and a shrunken keratin core (**Fig. 1G, Fig. S4**). Aggravatingly, DKO caused the loss of keratin core and the emergence of a Krt5^+^ Krt13^+^ cell population at the center of esophageal organoids (**Fig. S4D-E**). These results collectively demonstrate that either *Trp53*/*Cdkn2a*-dual loss or, to a lesser extent, *Trp53-single* loss induce neoplastic evolution that exhibits hyperproliferation and dysplasia across mouse oral/esophageal organoid models.

To extend the findings from murine organoid models to the human context, we next established genetically engineered human tongue organoid lines. While Ctrl human tongue organoids propagated for only 4–5 months, SKO and DKO modifications enabled their continuous expansion and propagation for over 21 months. Although SKO did not significantly affect organoid size or viability **(Fig. S2D-F**), it resulted in prominently enlarged nuclei, upregulated Ki67 labeling index, expanded Krt5+ basal cells, and decreased Krt13+ suprabasal cells, indicating the emergence of dysplastic features (**Fig. 1E-G**). Importantly, DKO significantly exacerbated all of these dysplastic features (**Fig. 1E-G**, **Fig. S2D-F**). These findings demonstrate highly consistent results across murine and human tissues, underscoring the robustness and rigor of these organoid models in faithfully capturing key aspects of dysplastic transformation.

### Additional Genetic Drivers Promote Full Malignant Transformation of Premalignant Organoids

Building on the CRISPR-edited organoid models, we next aimed to investigate the transition from precursor lesions to a malignant state. As outlined earlier, concurrent loss of *TP53* and *CDKN2A* is the most frequent genomic alteration in UASCC patients, and DKO organoids exhibit pronounced dysplastic features suggesting that they are primed for full malignant transformation. Thus, DKO organoids were selected as the foundational model for subsequent genomic editing. We targeted frequently mutated drivers in UASCC, including *PIK3CA*, *NOTCH1*, and *KMT2C*, which are known to occur against a backdrop of *TP53*/*CDKN2A* dual loss (**Fig. 2A**). Notably, these three genes are altered in a mutually exclusive manner in UASCC, suggesting they confer distinct biological properties and providing a strong rationale for their individual genetic engineering.

**Figure 2.**
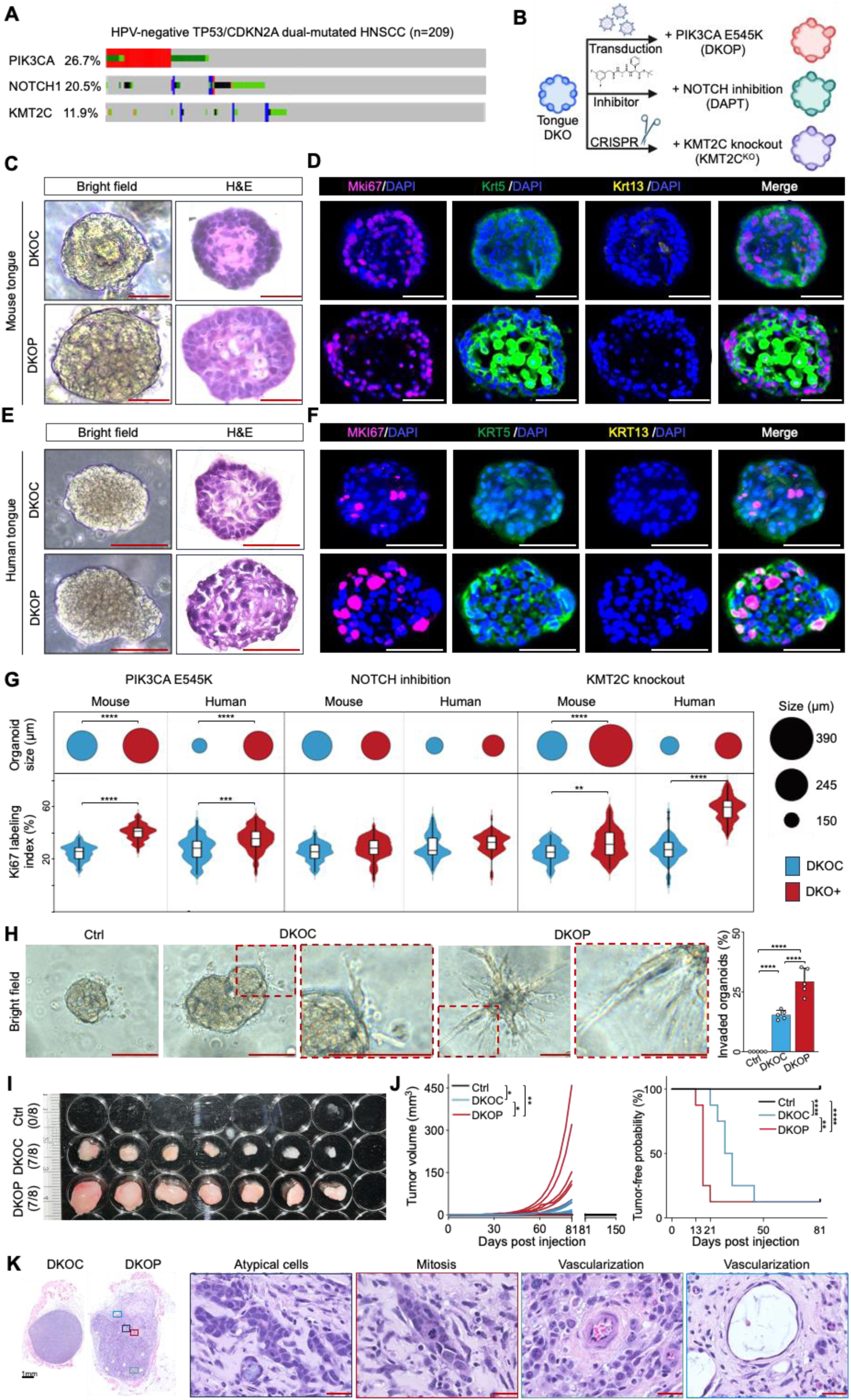
Additional genetic drivers promote full malignant transformation of premalignant organoids. **(A)** Genomic aberrations of *PIK3CA*, *NOTCH1*, and *KMTC2* in *TP53*/*CDKN2A* dual-mutated TCGA HNSCC samples, plotting via the Cbioportal (22). **(B)** Schematic of modeling the three common genomic aberrations in *TP53*/*CDKN2A* dual-mutated samples using mouse and human tongue DKO organoid models. This plot was created via Biorender. **(C)** Representative bright-field, H&E, and **(D)** IF images showing staining of Mki67 (magenta), Krt5 (green), and Krt13 (yellow) in 3-week-old mouse tongue DKOC and DKOP organoids. **(E)** Representative bright-field, H&E, and **(F)** IF images showing staining of MKI67 (magenta), KRT5 (green), and KRT13 (yellow) in 3-week-old human tongue DKOC and DKOP organoids. **(G)** Quantification of organoid size and Ki67 labeling index of 3-week-old mouse and human tongue DKO organoid lines with *PIK3CA^E545K^* over-expression, NOTCH inhibition, and *KMT2C* knockout (*n* = 50 per group). **(H)** Organoid invasion assay of mouse tongue Ctrl, DKOC, and DKOP organoids. Representative bright-field images of 10-day-old organoids are shown. The invasiveness of organoids cultured in a 3D matrix was quantified in the right bar graph based on the formation and enrichment of spindle-like protrusions (n = 5 independent experiments). **(I)** Allograft images from nu/nu mice subcutaneously injected with mouse tongue Ctrl, DKOC, and DKOP organoid cells (n=8 per group). **(J)** Comparison of tumor volume (left) and tumor-free survival (right). **(K)** Representative H&E images of DKOC and DKOP allografts. Scale bar, 100 μm. \**P* < 0.05; \*\**P* < 0.01; \*\*\**P* < 0.001; \*\*\*\**P* < 0.0001.

To model the gain-of-function mutations and amplifications of *PIK3CA*, we overexpressed one of its most frequent hotspot mutations in UASCC, *E545K.* We transduced *PIK3CA^E545K^* and corresponding control plasmid (see Methods section) into DKO organoids, generating DKO+*PIK3CA^E545K^* (termed DKOP) and DKO+control vector (termed DKOC) organoid lines (**Fig.2B**). IF staining verified the overexpression of *PIK3CA^E545K^* (**Fig. S5A**). Importantly, the introduction of *PIK3CA^E545K^* led to a significant increase in organoid size, enhanced Krt5 expression, and increased proliferation, accompanied by a loss of differentiation in both mouse and human organoid models (**Fig. 2C-G, Fig S5**).

Next, we modeled the loss of *NOTCH1* using the specific small-molecule inhibitor DAPT, as described in previous studies (23,24). The inhibition of NOTCH1 signaling was validated by a significant downregulation of canonical NOTCH target genes (**Fig. S6A**). Interestingly, while NOTCH inhibition had no additional effects on organoid growth in either mouse or human tongue DKO models, it significantly altered differentiation, evidenced by a marked increase in Krt5^+^Krt13^+^ cells in the inner part of organoids, a double-positive population uncommon in other models (**Fig. S6**). To model loss-of-function mutations and deletions of *KMT2C*, we employed CRISPR/Cas9 gene editing (**Fig. S7**), with successful targeting confirmed via Sanger sequencing (**Fig. S7A, G**). The loss of *KMT2C* substantially increased organoid size, proliferation, and the Ki67 labeling index (**Fig. S7C-F**) in both human and mouse DKO lines. Intriguingly, *Kmt2c* loss also induced striking morphological changes in murine organoids, shifting from solid squamous structures to cystic formations composed of either a single layer or stratified layers of well-organized cells (**Fig. S7E-F**). However, these morphological changes appeared to be specific to the murine models.

Although NOTCH inhibition and *KMT2C* loss each elicited neoplastic changes in DKO organoids, the *PIK3CA^E545K^* model demonstrated the most potent and consistent effects across both human and murine tissues. Consequently, we focused on the DKOC and DKOP models for further analyses. We first evaluated their in vitro invasiveness using a 3D matrix-based organoid invasion assay. Notably, compared to Ctrl organoids, DKOC exhibited significantly increased spindle-like protrusions, which were further elevated in the DKOP group, indicating enhanced invasive capacity (**Fig. 2H**). Next, we assessed the in vivo tumorigenic potential, the gold standard for assessing malignant transformation. Following subcutaneously injection into Nu/Nu mice, no tumor was observed upon the inoculation of Ctrl organoids, as expected. While both DKOC and DKOP groups formed 7 tumors out of 8 injections, DKOP grew considerably faster and resulted in significantly shorter tumor-free survival (**Fig. 2I-J**). Histological examination showed that DKOC grew into circumscribed, well-contained SCC (**Fig. 2K, Fig. S8**). In sharp contrast, DKOP tumors appeared more invasive, disorganized, poorly differentiated squamous cancer, closely resembling aggressive human HNSCC in histology and pathology. Indeed, malignant features such as atypical tumor cells, mitosis, and vascularization were prominent in DKOP but absent in DKOC tumors (**Fig.2K, Fig. S8**). These findings suggest that *PIK3CA^E545K^* fully transforms premalignant organoids into aggressive cancer. Furthermore, these results underscore our robust and valid organoid platform to model premalignant biology and early tumorigenesis, enabling rigorous investigation of underlying biological and molecular changes during squamous neoplastic transformation.

### Single-Cell Transcriptomic Analysis of Genetically-Engineered Organoid Models

As shown above, DKOC and DKOP organoids faithfully recapitulated the sequential squamous neoplastic evolution. To gain mechanistic insights into this malignant transformation process at single-cell resolution, we performed single-cell RNA-sequencing (scRNA-seq) on mouse tongue wildtype (Ctrl, 2616 cells), DKOC (2658 cells) and DKOP (1705 cells) organoids ( **Fig. 3A**). Uniform Manifold Approximation and Projection (UMAP) showed that DKOC/DKOP cells were largely separated from wildtype control cells (**Fig. 3A**), suggesting profound transcriptional perturbation elicited by the loss of *Trp53*/*Cdkn2a*. Four major clusters were identified and annotated based on established oral squamous cell markers (**Fig. 3B-C, Fig. S9**), including oral quiescent progenitors (e.g., Col17a1, Dst), proliferative basal cells (e.g., Krt14, Mki67, Top2a), early suprabasal cells (e.g., Krt4, Mal), and differentiated squamous cells (e.g., Sprr1a, Klf4). Importantly, compared with control organoids which were dominated by suprabasal and differentiated squamous cells, both DKOC and DKOP groups exhibited dramatically increased proliferative basal cells, accompanied by the loss of suprabasal and differentiated cells (**Fig. 3D**). Immunofluorescence staining of differentiated squamous cell marker Klf4 and proliferative basal cell marker Krt14 confirmed this strong phenotypic shift (**Fig. 3E**), which is also consistent with our earlier observations (**Figs. 1**-**2**).

**Figure 3.**
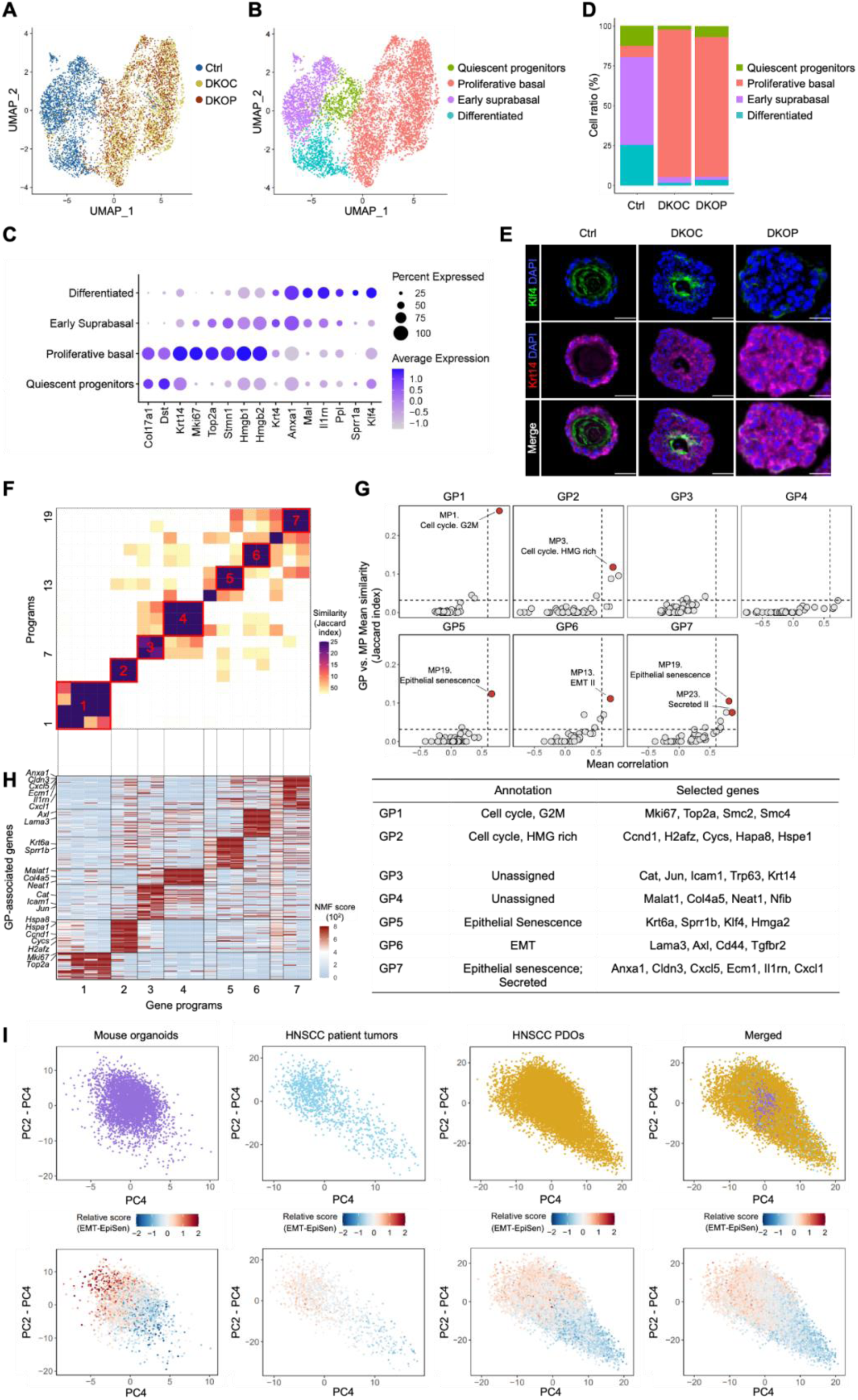
ScRNA-seq analysis of genetically-engineered organoid models. **(A)** UMAP projecting single-cell profiles of mouse tongue Ctrl, DKOC, and DKOP organoids and **(B)** cluster annotations. **(C)** Dot plot of cell marker genes for each cluster. **(D)** Bar chart showing the proportion of cell clusters across each organoid line. **(E)** IF staining images for the differentiation marker Klf4 (green) and proliferative basal marker Krt14 (magenta) in mouse tongue organoid lines. **(F)** Heatmap of unsupervised clustering of NMF programs based on program genes, with similarity measured by Jaccard indices. Seven clusters of consensus gene programs (GPs) are numbered and highlighted in red frames. **(G)** Dot plot showing the mean Jaccard index (y axis) between the murine GPs and published meta-programs (29), and mean correlation with single-cell scores (x axis). Dashed lines indicate a 99.9% confidence threshold determined by permutations. **(H)** Left: heatmap of the relative expression of GP genes, with each GP aligned to **(F)**. Right: annotation and selected top genes for each of the seven GPs. **(I)** PCA plots of indicated single-cell samples. Upper panel: Cells are colored by sample origin (mouse organoids, HNSCC patient tumors, or HNSCC PDOs). Lower panel: Cells are colored by relative scores for EMT and EpiSen GP genes identified from mouse organoids. Scale bar, 100 μm.

To understand molecular mechanisms underlying this neoplastic evolution process, we focused on characterizing gene modules or programs, defined as sets of coordinately expressed genes, that are frequently associated with cellular states and processes (25–28). We applied the non-negative matrix factorization (NMF) method to each organoid line to identify robust gene programs (GPs), as described previously (27–30). We clustered the NMF programs by their shared fractions of top genes (quantified by Jaccard indices), which identified seven GPs clusters (**Fig. 3F**). We subsequently defined a consensus GP from each of these 7 clusters based on shared genes in at least two organoid lines (**Fig. 3F**).

To understand these seven consensus GPs, we annotated them by performing enrichment analysis of their signature genes against well-curated meta-programs (MPs) established by the NMF approach from large-scale scRNA-seq delineation of pan-normal/tumor samples (29). Five of the seven GPs showed significant similarity (false discovery rate (FDR) q < 10^−9^; hypergeometric test) and high correlation of single-cell scores (**Fig. 3G**) with previously defined MPs (29). Specifically, these GPs were annotated to Cell cycle (GP1/2), Epithelial-mesenchymal transition (EMT, GP6), and Epithelial senescence (EpiSen, GP5/7).

To determine whether the GPs identified in our mouse organoids effectively recapitulate those in HNSCC patients, we performed an integrated analysis of our mouse organoid data with scRNA-seq data from in vivo HNSCC patient tumor samples from Puram et al. (24) and in vitro HNSCC patient-derived organoids (PDOs) from our recent study (10). Indeed, PCA plots revealed that mouse organoid cells, HNSCC patient tumor cells and PDOs were intermingled, overlapping transcriptional space (**Fig. 3I**, upper panel) and indicating substantial transcriptional comparability between the models. As expected, expression heterogeneity was lower in mouse organoids, reflecting their controlled genetic context with early neoplastic cells driven by a limited number of mutations. In contrast, patient tumors and PDOs exhibited greater heterogeneity due to their diverse mutational landscapes, distinct driver combinations, and accumulation of copy number alterations over time.

We next selected EpiSen and EMT programs from our mouse organoid models and plotted relative GP scores for each single cell. These GPs were chosen as they were reported to represent two distinct, polarized cellular states within subsets of intratumoral malignant cells (31). Consistent with this notion, in our murine organoid samples, EMT+ and EpiSen+ cells were distinctly separated, occupying two extreme regions (**Fig. 3I**, bottom panel). Importantly, our murine organoid GP genes also successfully segregated and identified EMT+ and EpiSen+ cells in both HNSCC patient tumors and PDOs, albeit to a lesser extent (**Fig. 3I**, bottom panel).

We further conducted a reciprocal analysis by applying human HNSCC tumor GP genes to our mouse organoid cells and plotting their relative GP scores. Importantly, we confirmed that human patient GP genes effectively separated EMT+ and EpiSen+ states in the murine cells (**Fig. S10A**). This result was further reproduced using the GP genes from HNSCC PDOs (**Fig. S10B**). Collectively, these findings suggest that the GPs identified in our genetically-engineered murine organoids faithfully recapitulate the cellular states and transcriptional heterogeneity observed in human tumors, both in vivo and in vitro.

### Prominent Downregulation of a Senescence Program During Oral Neoplastic Evolution

Next, we mapped each GP to individual single cells and compared the ratios of each GP across the three organoid groups (**Fig. 4A-B**). Consistent with our phenotypic data (**Figs. 1**-**2**), cell cycle programs (GP1 and GP2) showed a substantial increase from Ctrl to DKOC/DKOP organoids. Intriguingly, EpiSen programs (GP5 and GP7) demonstrated marked downregulation (**Fig. 4B**). Given that senescence-associated secretory phenotype (SASP) is a hallmark of cellular senescence (32), we were particularly interested in GP7, which was also enriched in the Secreted program (**Fig. 3G-H**). Moreover, GP7 showed progressive downregulation during neoplastic evolution.

**Figure 4.**
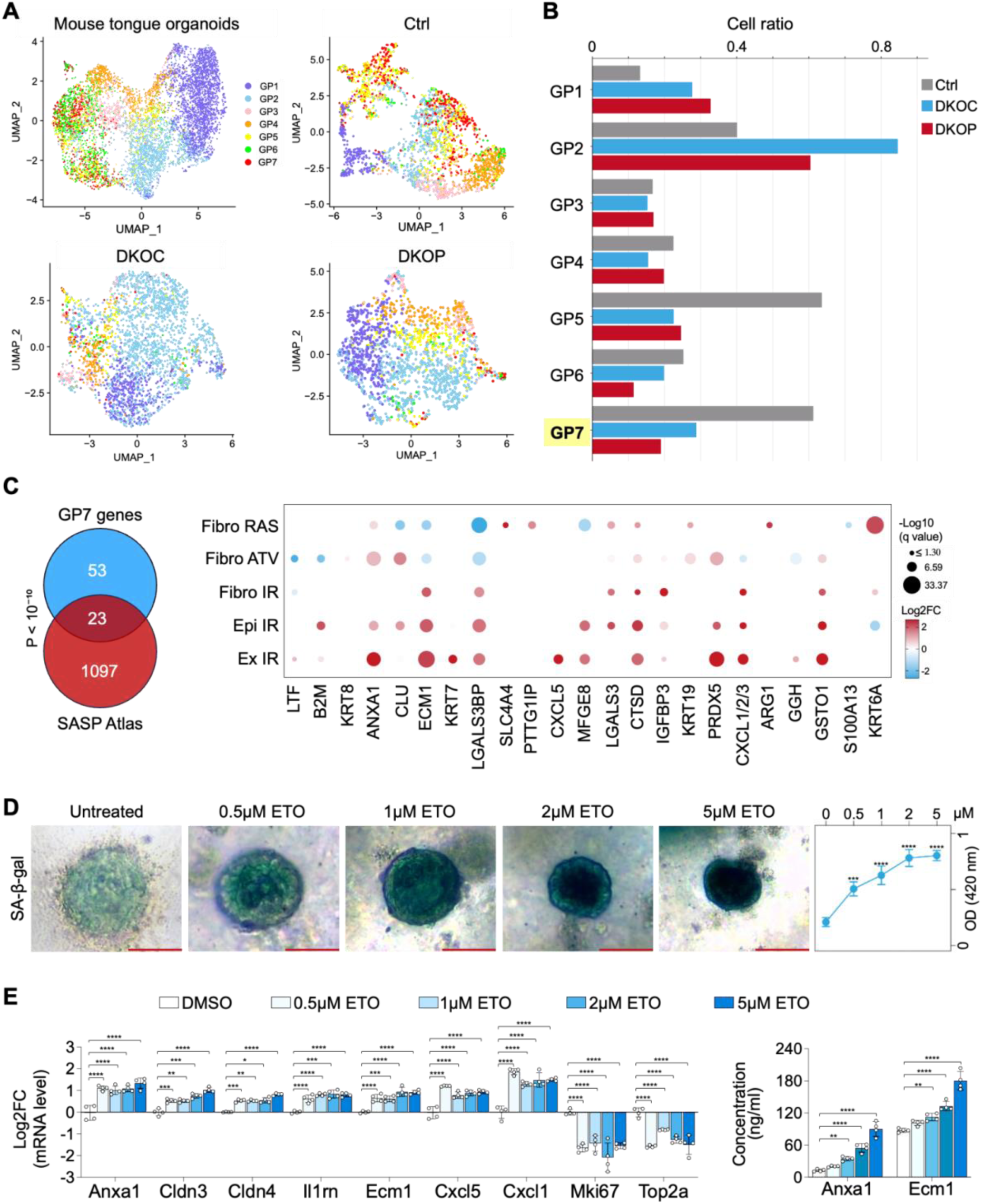
Significant downregulation of a senescence program during oral squamous neoplastic evolution. **(A)** UMAPs of the distribution and **(B)** cell ratio of each gene program (GP) at the single-cell level in mouse tongue organoid models. **(C)** Venn diagram illustrating the overlap between genes identified in GP7 and the senescence-associated secretory phenotype (SASP) Atlas dataset (29). Fibro RAS: RAS overexpression-induced senescence in fibroblasts; Fibro ATV: atazanavir treatment-induced senescence in fibroblasts; Fibro IR: irradiation-induced senescence in fibroblasts; Epi IR: irradiation-induced senescence in epithelial cells; Ex IR: exosomal SASP from senescent fibroblasts post-irradiation. **(D)** Induction of senescence in mouse tongue Ctrl organoids treated with etoposide (ETO), confirmed by SA-β-gal staining (*n* = 3 per group). **(E)** Left: relative mRNA expression levels of representative GP7 genes and cell cycle genes in ETO-treated mouse tongue Ctrl organoids. Right: protein concentrations of secreted Anxa1 and Ecm1 from ETO-treated mouse tongue Ctrl organoids. Scale bar, 100 μm. \**P* < 0.05; \*\**P* < 0.01; \*\*\**P* < 0.001; \*\*\*\**P* < 0.0001.

To characterize the GP7 program, we first examined its program genes (n=76) for their association with SASP using the SASP Atlas database (33), a proteomic database of soluble proteins and SASP factors originating from multiple different senescent inducers. Importantly, 23 out of these 76 factors exhibited significantly overlapped protein secretion in the SASP secretome upon senescence induced by various stresses (e.g., IR-radiation, oncogene activation by RAS) (*P* < 10^-10^, hypergeometric test; **Fig. 4C**).

To further validate the association of GP7 genes with cellular senescence, we treated murine tongue organoids with Etoposide, a chemotherapeutic agent known to induce senescence (34). The induction of the senescence phenotype was validated by senescence-associated β-galactosidase (SA-β-gal) staining (**Fig. 4D**) and potent downregulation of cell cycle genes, such as Mki67 and Top2a (**Fig. 4E**). Importantly, representative GP7 genes (those exhibiting > 2 Log2FC and overlapping with the SASP Atlas and/or the human HNSCC EpiSen program, see Methods) were consistently upregulated following Etoposide treatment (**Fig. 4E**). Additionally, increased protein secretion of select SASP factors, including Anxa1 and Ecm1, was validated using ELISA assays (**Fig. 4E**). These findings strongly suggest that GP7 is linked to cellular senescence and its progressive downregulation corresponds to the stepwise loss of senescence-associated pathways during early neoplastic evolution.

### ANXA1 Is a Critical Regulator of the Senescence Program and Tumor Suppressor in Early Squamous Neoplastic Progression

Captivated by the stepwise downregulation of the senescence program during oral neoplastic evolution, we next sought to identify its functional regulator(s). We performed a correlation analysis between the senescence program score and the expression of its 76 program genes (**Fig. 5A**). Among these, Anxa1 emerged as a prominent candidate: its expression exhibited a strong correlation with the senescence program score (**Fig. 5A**) and was downregulated from control to DKOC/DKOP organoids in the murine scRNA-seq (**Fig. S11A**). Moreover, Anxa1 itself is a component of the SASP secretome and was significantly over-secreted in various senescent contexts (**Fig. 4C, E**).

**Figure 5.**
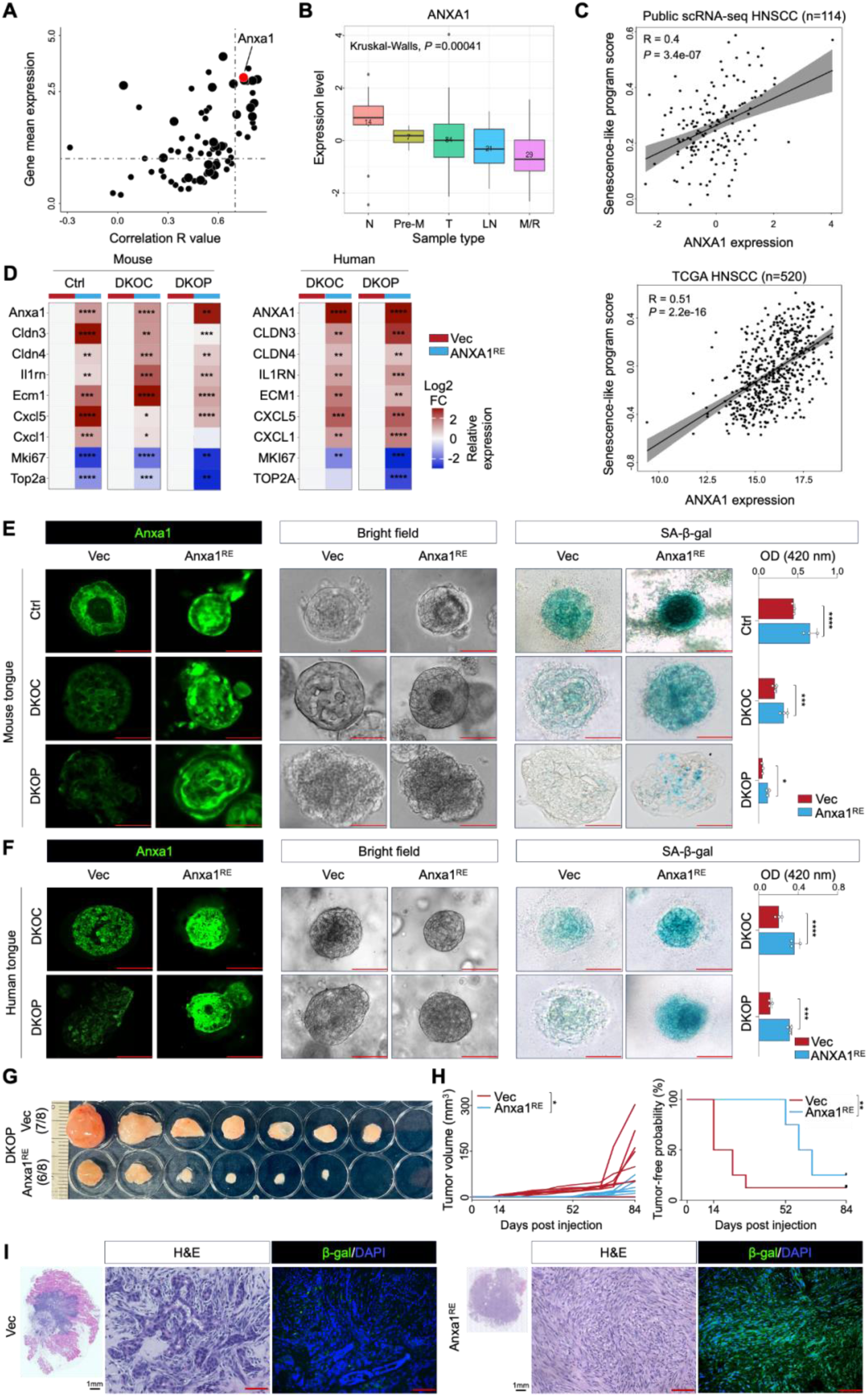
ANXA1 is a critical regulator of the senescence program. **(A)** Scatterplot showing the correlation (*R* value) between GP7 genes and their expression levels in control organoids from scRNA-seq data. Large-sized and small-sized dots denote genes expressed in all and <half of organoids, respectively. **(B)** Epithelial-specific expression of ANXA1 across distal normal samples (N), premalignant lesions (Pre-M), primary tumors (T), lymph node samples (LN), and metastasis/recurrent tumors (M/R) from an integrated human cohort of HNSCC scRNA-seq data. **(C)** Upper panel: correlation between ANXA1 expression and the GP7 score in human HNSCC scRNA-seq. Lower panel: correlation between ANXA1 expression and the GP7 score in TCGA HNSCC samples. **(D)** Relative mRNA expression levels of representative GP7 genes and cell cycle genes in mouse and human tongue organoids transduced with either control vector (Vec) or ANXA1-restoration vector (ANXA1^RE^) (*n* = 3 per group). **(E-F)** Representative images of IF staining for Anxa1, bright-field, and SA-β-gal staining in Vec and ANXA1^RE^ mouse **(E)** or human **(F)** tongue organoids. **(G)** Allograft images from nude mice subcutaneously injected with mouse tongue Vec and Anxa1^RE^ DKOP organoid cells (*n* = 8 per group). **(H)** Comparison of tumor volume (left panel) and tumor-free survival (right panel) in allograft experiments. **(I)** Representative H&E images and IF staining for β-Galactosidase in Vec and Anxa1^RE^ DKOP allografts. Scale bar, 100 μm. \**P* < 0.05; \*\**P* < 0.01; \*\*\**P* < 0.001; \*\*\*\**P* < 0.0001.

To validate these findings in human tissues, we analyzed epithelial-specific ANXA1 expression in a combined scRNA-seq cohort of public (3,28,35,36) and internal HNSCC patient samples (n=114). Remarkably, this independent dataset revealed a significant, progressive downregulation of ANXA1 from distal normal tissues, through premalignant lesions, primary tumors, lymph node metastases, to recurrent/metastatic tumors (**Fig. 5B**). Furthermore, ANXA1 expression was consistently correlated with the senescence program score across this independent human scRNA-seq dataset (**Fig. 5C**, upper panel). In addition, TCGA HNSCC bulk RNA-seq data confirmed the downregulation of ANXA1 in tumor samples, and its expression was again positively correlated with the senescence program score (**Fig. 5C**, lower panel). These findings position ANXA1 as a potential functional regulator of the senescence program, with its loss being closely linked to neoplastic progression in HNSCC.

We next investigated the biological significance of ANXA1 by ectopically restoring its expression in organoid lines. In the DKOC/DKOP models, Anxa1 expression was reduced 1.8- to 1.9-fold compared to controls (**Fig. S11A**). To counteract this downregulation, we carefully titrated ANXA1 ectopic expression to achieve a 1.5- to 3-fold increase of its mRNA levels (**Fig.S11B**). Notably, restoration of ANXA1 expression significantly upregulated the expression of senescence program genes and concomitantly suppressed cell cycle gene expression across various genotypes of both human and murine organoid models (**Fig. 5D**). Functionally, ANXA1 restoration robustly enhanced cellular senescence, as evidenced by increased SA-β-gal staining, and markedly reduced organoid growth and viability across both murine and human organoid groups (**Fig. 5E–F, Fig. S11C-D**). These data demonstrate that ANXA1 regulates the senescence program, potentially contributing to the inhibition of oral malignant transformation.

To evaluate whether Anxa1 could inhibit neoplastic progression in vivo, we subcutaneously injected nude mice with DKOP organoids transfected with either control vector (Vec) or their Anxa1-restored counterparts (Anxa1^RE^). Remarkably, Anxa1 restoration resulted in significantly smaller tumors, markedly increased tumor-free survival, and extended latency compared with the control group. Indeed, the first tumor in the Anxa1^RE^ group appeared at day 52 post-injection, in stark contrast to day 14 in the control group (**Fig. 5G-H**). Crucially, histological analysis revealed that Anxa1^RE^ tumors exhibited reduced atypia, decreased cellular pleomorphism, lower mitotic activity, and diminished cellularity, resulting in a lower tumor grade. Some regions of Anxa1^RE^ tumors even displayed nearly normal histomorphology (**Fig. 5I, Fig. S12**). Furthermore, enhanced β-gal staining was observed in the Anxa1^RE^ group (**Fig. 5I**). Collectively, these results establish ANXA1 as a critical regulator of the senescence program and highlight its tumor-suppressive role in oral squamous neoplastic progression. This represents a novel and significant mechanism by which ANXA1 inhibits tumorigenesis at the earliest stages of cancer development.

### A Novel ANXA1-SMAD3-p27^KIP1^ Pathway Regulating Cellular Senescence during Early Neoplastic Evolution

In exploring the mechanisms underlying the regulation of ANXA1 on cellular senescence, we first considered canonical regulators of this process: TP53 (37), CDKN2A (38), CDKN1A (p21^CIP1^) (39), RB (40), CDKN1B (p27^KIP1^) (41,42). Since both *TP53* and *CDKN2A* were null in our DKOC/DKOP model, we assessed the expression of the remaining three factors. Notably, ANXA1 promoted the expression of p27^KIP1^ across both murine and human organoid models (**Fig. 6A**), while p21^CIP1^ or RB remained unchanged (**Fig. S13**). The upregulation of p27^KIP1^ was further verified at the protein level (**Fig. 6B**). We then focused on p27^KIP1^ and performed rescue assays. Importantly, silencing of p27^KIP1^ reversed ANXA1-induced senescence as measured by both senescence-program gene expression (**Fig. 6C**) and SA-β-gal staining (**Fig. 6D**). Consistently, knockdown of p27^KIP1^ also restored organoid viability and growth which were inhibited by ANXA1 (**Fig. 6E**). These epistatic data suggest that p27^KIP1^ acts as a key downstream mediator of ANXA1-regulated cell senescence and early neoplastic evolution.

**Figure 6.**
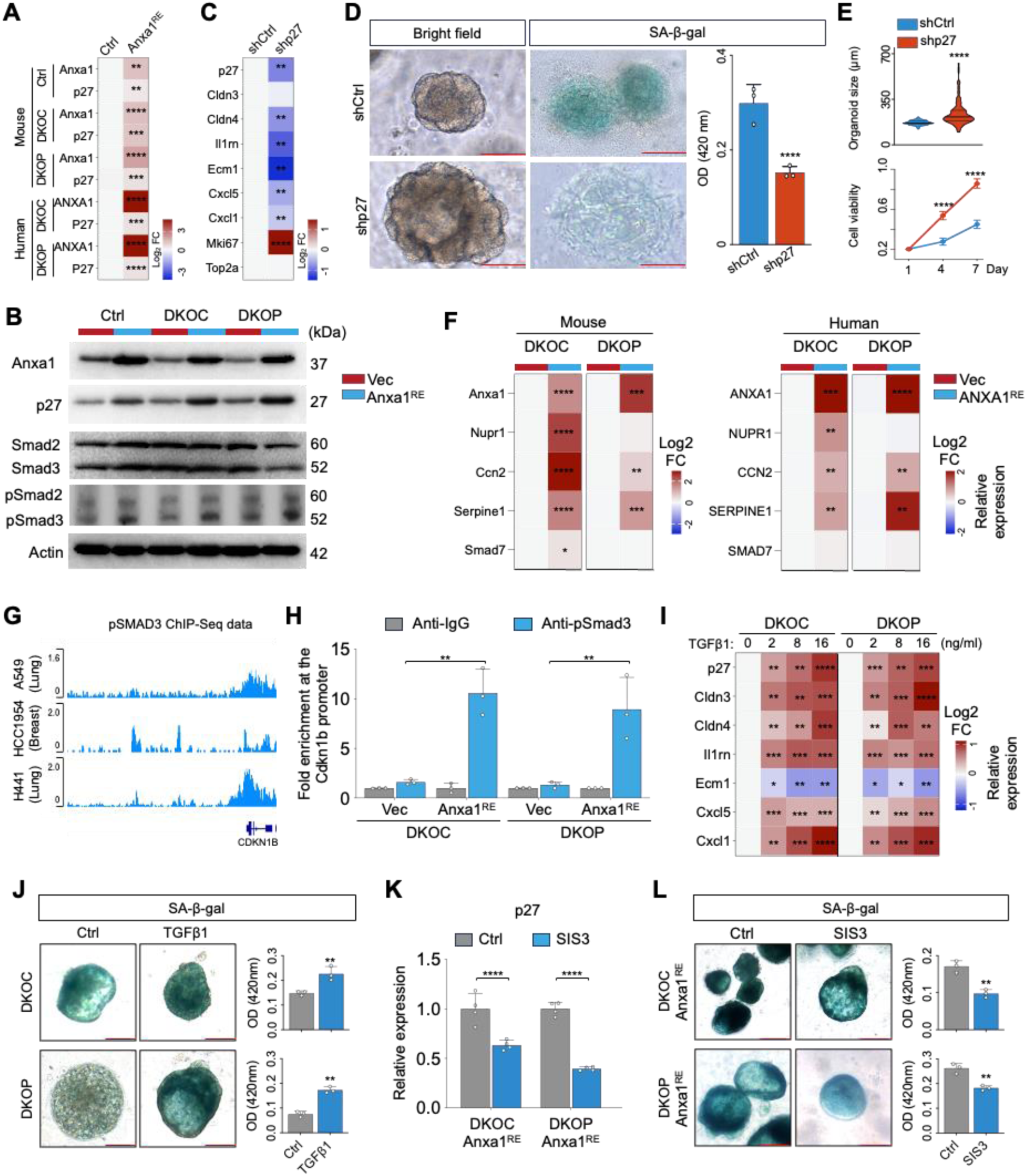
A novel ANXA1-SMAD3-p27^KIP1^ pathway regulating cellular senescence during early neoplastic evolution. **(A)** Relative mRNA expression levels of Anxa1 and p27 in mouse and human tongue organoids transduced with either control vector (Vec) or ANXA1-restoration vector (ANXA1^RE^). **(B)** Immunoblot detection of Anxa1, p27, Smad2/3, and pSmad2/3 in mouse tongue Vec and ANXA1^RE^ organoids. **(C)** Relative mRNA expression levels of representative GP7 genes and cell cycle genes in shCtrl and shp27 mouse tongue Anxa1^RE^ DKOP organoids. **(D)** Representative bright-field and SA-β-gal staining images of shCtrl and shp27 mouse Anxa1^RE^ DKOP organoids. **(E)** Organoid size (*n* = 50) and viability of shCtrl and shp27 mouse Anxa1^RE^ DKOP organoids, as assessed by the WST-1 assay (*n* = 6). **(F)** Relative mRNA expression levels of established SMAD3 direct target genes in mouse and human tongue Vec and ANXA1^RE^ DKOC and DKOP organoids (*n* = 3). **(G)** Public ChIP-Seq data showing pSMAD3 binding at the *CDKN1B* promoter locus. **(H)** ChIP-qPCR analysis showing the fold enrichment of pSmad3 occupancy at the Cdkn1b promoter region relative to IgG. **(I)** Relative mRNA expression levels of p27 and representative GP7 genes in mouse tongue DKOC and DKOP organoids treated with TGFβ1 for 48 hours (*n* = 3). **(J)** SA-β-gal staining of mouse tongue DKOC and DKOP organoids treated with TGFβ1 for 48 hours. **(K)** Relative mRNA expression levels of p27 in Anxa1^RE^ mouse DKOC and DKOP organoids treated with a specific inhibitor of pSMAD3 (SIS3) for 48 hours. **(L)** SA-β-gal staining of mouse Anxa1^RE^ DKOC and DKOP organoids treated with SIS3 for 48 hours. Scale bar, 100 μm. \**P* < 0.05; \*\**P* < 0.01; \*\*\**P* < 0.001; \*\*\*\**P* < 0.0001.

In the above results, upregulation of p27^KIP1^ occurred at the transcriptional level; since ANXA1 is not a transcription factor, this suggests an indirect regulatory mechanism. Thus, we investigated the transcription factors known to bind directly to and activate the transcription of p27^KIP1^, including FOXO1 (43), FOXO3a (44), SMAD ^(45)^, E2F1 (46). While TP53 is also a major transcriptional activator of p27^KIP1^, it is absent in our DKOC/DKOP models. Interestingly, we found that pSmad3 was consistently upregulated in ANXA1-restored organoids (**Fig. 6B**), accompanied by increased expression of established SMAD3 direct target genes, including NUPR (47), CCN2 (48), SERPINE (49), and SMAD7 (50) (**Fig. 6F**), while the remaining candidates showed no significant change in both human and murine organoids (**Fig. S13**). Moreover, analysis of public ChIP-Seq data confirmed that pSMAD3 directly bound to the *Cdkn1b* locus across various cell types, such as lung and breast cells (**Fig. 6G**). We next performed pSmad3 ChIP-qPCR in our oral organoid models and validated that endogenous pSmad3 showed strong occupancy at the *Cdkn1b* promoter. Importantly, Anxa1 restoration led to a 6.6-fold increase in pSmad3 enrichment at the *Cdkn1b* promoter compared to control, across both murine and human organoids (**Fig. 6H**).

Since SMAD3 phosphorylation and activity are primarily regulated by the TGFβ signaling, we stimulated our organoid models with the TGFβ1 protein, which consistently upregulated p27^KIP1^ transcription, enhanced senescence-program gene expression and induced cellular senescence (**Fig. 6I, J**), thereby phenocopying the effects of Anxa1 restoration. Conversely, the specific inhibitor of SMAD3 (SIS3) (51), which attenuates the phosphorylation of Smad3, diminished the expression of p27^KIP1^ and reduced senescence in Anxa1-restored organoids (**Fig. 6K, L**). Together, these data identify a novel ANXA1-SMAD3-p27^KIP1^ signaling axis that regulates cellular senescence during early neoplastic evolution.

### Genetically-defined organoid models for drug screening and validation

Despite extensive genomic characterization of UASCC, the translation of these findings into clinical practice remains limited. Currently, apart from immunotherapy, the only FDA-approved targeted therapy for UASCC is Cetuximab (specifically for HNSCC), an anti-EGFR antibody. However, Cetuximab has shown limited efficacy, and its use is in fact not guided by genetic testing (52), thus falling short of being considered a true genome-guided treatment. This underscores a critical gap in HNSCC management: the need for systematic, unbiased identification of drug-gene associations that are both clinically relevant and actionable.

In this context, our genetically-defined murine organoid models are uniquely positioned to identify drugs specifically effective against mutant *PIK3CA*. To enhance clinical relevance, we have concurrently generated a cohort of HNSCC PDO lines, with or without PIK3CA-activating alterations, identified by whole-exome sequencing (WES, **Fig.7G**). To further facilitate this drug discovery effort, we have developed innovative technologies to generate bioprinted 3D organoids according to a geometry that facilitates rapid and automated high throughput drug screening, integrated with label-free, time-resolved imaging analyzed using machine learning at single-organoid resolution. We have demonstrated that this advanced platform enables high-throughput, unbiased identification of either transient or persistent drug responsiveness across diverse tumor types (53–55).

**Figure 7.**
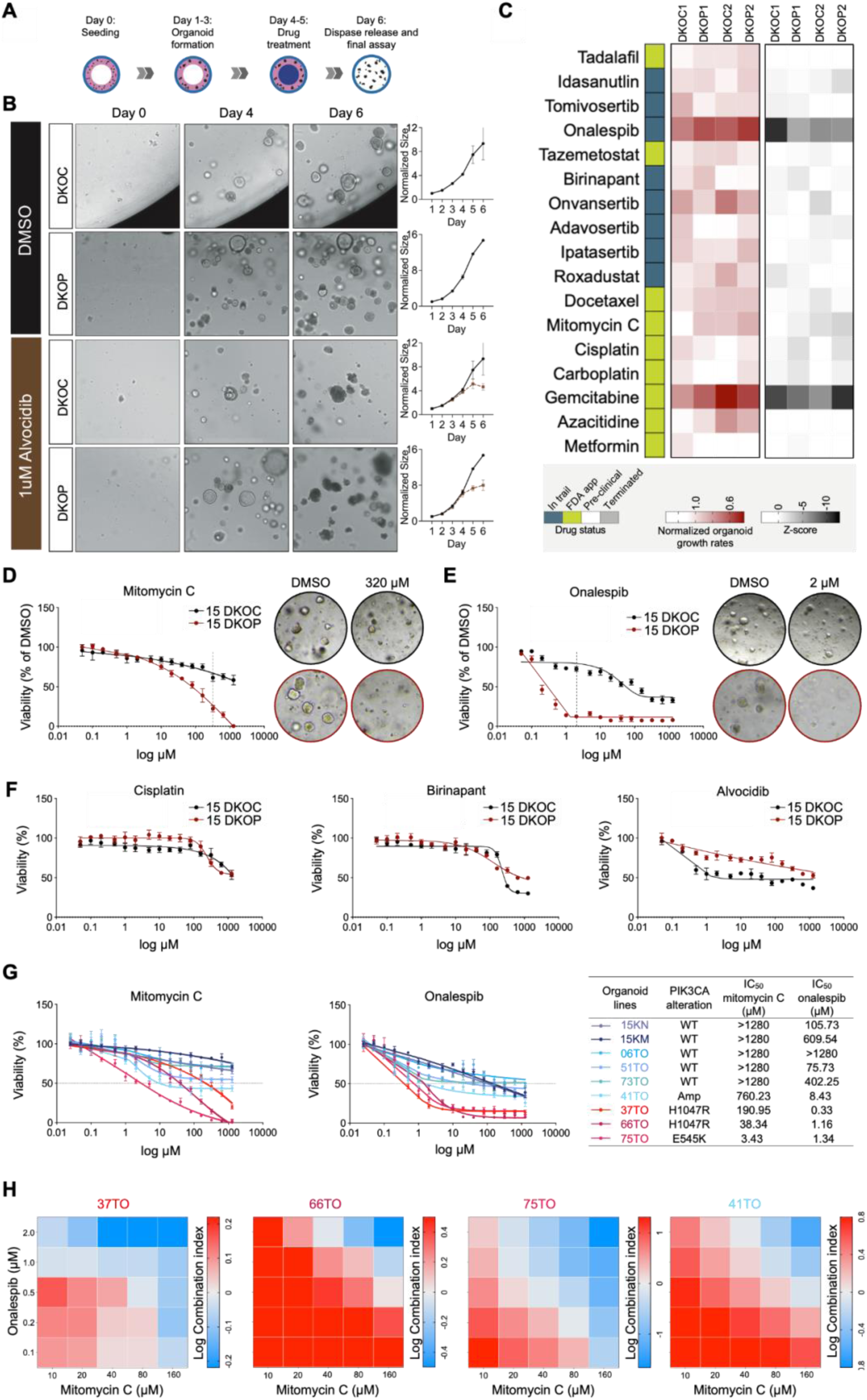
Genetically-defined organoid models for drug screening and validation. **(A)** Schematic of the experimental setup for drug screening. **(B)** Representative bright-field images and growth curves of organoids analyzed using machine learning-based segmentation. **(C)** Heatmaps of growth rates and ATP assay results (z-scores) for organoids. Growth rates were calculated as the ratio of organoid growth on day 6 (prior to the ATP assay) to day 4 (before the initial drug treatment), normalized to the DMSO vehicle control. Data from each of the two independent biological replicates are shown. **(D-F)** Sensitivity of human DKOC and DKOP organoid lines to five candidate drugs identified from the initial mouse organoid screening (n=3). **(G)** Sensitivity analysis of genetically-engineered human organoid lines and patient-derived HNSCC organoid lines with either wild-type or mutant/amplified PIK3CA, tested against Mitomycin C and Onalespib (n=3). **(H)** Combination index (CI) of Mitomycin C and Onalespib across *PIK3CA*-mutant (37TO, 66TO, and 75TO) and *PIK3CA*-amplified (41TO) PDOs (n=3). Log CI > 0 indicates antagonism; Log CI = 0 indicates additivity; Log CI < 0 indicates synergy. The data is normalized to DMSO vehicle control.

We thus applied this unbiased, single-organoid resolution screening platform to our DKO and DKOP models, initially testing a panel of 50 anti-cancer drugs, including FDA-approved agents and compounds in late-stage clinical trials (**Fig. 7A**). This system enables image-based quantification of individual organoids through machine learning-based segmentation (**Fig. 7B-C, Fig. S14A**). To validate the robustness of our approach, we concurrently performed an orthogonal, conventional ATP-based viability assay (**Fig. 7C, Fig. S14A**). Notably, this independent ATP-based measurement exhibited a strong correlation with organoid viability as assessed by machine learning-based image analysis (**Fig. S14B**). Moreover, in addition to technical replicates, we conducted two independent biological replicates using separately cultured organoid lines, which confirmed the high reproducibility of our finding.

Among the 50 drugs tested, five candidates (Alvocidib, Cisplatin, Birinapant, Mitomycin C, Onalespib) demonstrated greater inhibition of proliferation in DKOP organoids compared to DKOC organoids in at least one assay, with consistent trends observed across two trials (**Fig. 7C** and **Fig. S14A**). It was worth noting that PI3K-targeting inhibitors (Copanlisib, Alpelisib, Umbralisib, Duvelisib, Eganelisib) showed no significant difference in efficacy between DKOC and DKOP organoids (**Fig. S14C**), aligning with previous reports (56,57). These data underscore the complexity of gene-drug associations, as mutation-directed therapies frequently fail to produce the anticipated effects, reinforcing the necessity of unbiased screening approaches.

We then conducted a 15-point concentration counter-screening in human DKOC and DKOP lines for the five drugs identified in the initial mouse organoid screening. Validating the drug screen data, both Mitomycin C and Onalespib demonstrated significantly enhanced cytotoxicity in DKOP compared to DKOC organoids, while the other three drugs did not exhibit this selective effect (**Fig. 7D-F**). To confirm the *PIK3CA*-specific response to Mitomycin C and Onalespib, we expanded the validation to additional genetically-engineered human lines: DKOC with NOTCH inhibition (15KN) and DKOC with *KMT2C*^KO^ (15KM). We further included seven HNSCC patient PDO lines with or without *PIK3CA*-activating alterations (i.e., hotspot mutations/gene amplification) detected by WES (**Fig. 7G**, right panel). Importantly, *PIK3CA*-mutant/amplified lines showed markedly higher sensitivity to Mitomycin C and Onalespib treatment compared to *PIK3CA* wild-type counterparts (**Fig. 7G**).

Interestingly, while Mitomycin C caused a gradual and steady decline in cell viability, Onalespib elicited a sharp drop in viability at very low doses, with its effect plateauing at higher doses (**Fig. 7 D-E**,). These complementary drug response patterns suggest a potentially synergistic effect between these drugs. Indeed, compared to single treatments, the combination of low dose of Mitomycin C and Onalespib resulted in increased cell death in *PIK3CA*-mutant PDO lines (**Fig. 7H, Fig. S15**). However, this synergistic effect was not observed in *PIK3CA* wild-type PDOs (**Fig. S15**). These results together suggest that the combination of Mitomycin C and Onalespib may offer a promising therapeutic strategy specifically for *PIK3CA*-driven HNSCC, further underscoring our genetically-defined, cross-species organoid models as valid, clinically-relevant platforms for drug-gene association discovery.

## DISCUSSION

In this study, we leveraged genetically-engineered organoid models to dissect the early molecular and cellular events underlying malignant evolution of UASCC. We demonstrated the critical role of *TP53*/*CDKN2A* double knockout in initiating neoplastic transformation. This baseline was further amplified by additional oncogenic drivers such as *PIK3CA*, *NOTCH1* or *KMT2C*, highlighting the synergistic effects of cooperative mutations in driving malignant progression. Single-cell and computational analyses provided a detailed map of cellular changes during neoplastic evolution, revealing a marked loss of differentiated squamous cells accompanied by a concomitant expansion of quiescent basal and proliferative squamous cells. Importantly, these cellular transitions were accompanied by the progressive downregulation of a senescence program, which emerged as a hallmark of early squamous neoplasia.

Cellular senescence was initially recognized as a fundamental stress-response process designed to halt the proliferation of damaged or aberrant cells, thereby preventing the development of diseases such as cancer (58). However, recent compelling data has challenged this traditional view, suggesting that senescent cells may paradoxically contribute to cancer development and progression (59). For example, increasing evidence indicates that subsets of senescent cells can be released from growth arrest and re-enter the cell cycle, exhibiting enhanced cellular stemness and plasticity and contributing to tumor progression (60,61). Thus, our understanding of the intricate interplay between senescence and cancer remains incomplete. The precise mechanisms by which senescent cells influence cancer biology represent a critical knowledge gap. A major roadblock to investigating senescence in cancer is the lack of valid models that capture this dynamic process, as most senescence research relies on either fibroblasts or cell line models. Moreover, rigorous investigation of senescence during early malignant evolution is scarce.

Cellular senescence can be induced by a variety of stress stimuli, such as DNA damage, oncogene activation and metabolic stress (62). While TP53, RB, CDKN2A, CDKN1A/B are major regulators mediating these stress signals to induce senescence, other regulators (e.g., PTEN, CSN5 and TAp63) have also been identified (63–65). Here, our scRNA-seq data from murine organoids identified ANXA1 as a novel regulator of the senescence program, which is corroborated by both scRNA-seq and bulk RNA-seq data from human samples. ANXA1, a phospholipid-binding protein with immunomodulatory function, has been investigated in cancer (66). However, a notable knowledge gap exists concerning contradictory reports on the function of ANXA1, whether it acts as either a protumor or antitumor factor (67). These conflicting data suggest that the expression and function of ANXA1 may be regulated in a tissue- and tumor-specific manner. The inconsistency may also stem from the fact that almost all prior research has utilized cancer cell line models, whereas our approach uses exclusively organoid models. Importantly, we showed that ANXA1 functionally promoted cellular senescence and inhibited neoplastic features across multiple murine and human organoid models, both in vivo and in vitro. This data underscores ANXA1 as a crucial senescence regulator, acting as a biological guardrail to prevent malignant transformation.

Despite abundant genomic alterations in UASCC that are shared with other tumor types for which targeted therapies have been successfully developed, Cetuximab, an anti-EGFR antibody, remains the only approved gene-targeting drug for UASCC. However, its clinical utility is limited, with a response rate of only about 10%, leaving the majority of patients without effective targeted options. Compounding this challenge is the scarcity of drug screens specifically tailored to UASCC, highlighting a critical unmet need in UASCC management: the systematic and unbiased identification of clinically relevant drug-gene associations. Achieving this goal is impeded by two primary barriers: (i) the significant intertumoral heterogeneity driven by diverse genomic mutations, and (ii) the lack of robust human models capable of defining genotype-phenotype relationships in a rigorous and translationally relevant manner.

Our genetically-defined, cross-species organoid models are uniquely suited to overcome the barrier of intertumoral heterogeneity, enabling the systematic exploration of how specific genomic lesions influence drug responses. To further enhance the drug discovery pipeline, we integrated advanced technologies, including bioprinted 3D organoid platforms coupled with label-free, time-resolved imaging analyzed by machine learning at single-organoid resolution. Using this platform, we conducted an unbiased, high-throughput drug screen and identified a novel gene-drug association for *PIK3CA*-mutant HNSCC. The findings were further validated in patient-derived organoid models, highlighting Mitomycin C and Onalespib as selectively effective agents for *PIK3CA*-mutant tumors. This study not only demonstrates the translational potential of our organoid-based drug discovery pipeline but also provides a foundation for developing targeted therapeutic strategies tailored to the unique genetic landscape of UASCC. Such efforts are crucial for addressing the longstanding gap in precision medicine for this aggressive and heterogeneous cancer type.

While our study offers valuable insights into the mechanisms driving early squamous neoplastic evolution and potential therapeutic strategies, several limitations warrant discussion. Although the organoid models successfully recapitulate many aspects of human neoplasia, additional validation in in vivo systems is necessary to fully establish their relevance to patient tumors and the broader tumor microenvironment. Furthermore, the precise upstream regulatory mechanisms controlling ANXA1 expression during early tumorigenesis remain unclear, necessitating further investigation to better understand its role in early neoplastic suppression. Lastly, while Mitomycin C and Onalespib demonstrated efficacy in *PIK3CA*-mutant organoids, preclinical studies and animal experiments are essential to confirm their therapeutic potential and synergistic effects in vivo, in order to enhance translational relevance to clinical applications.

In summary, our study uncovers critical drivers and molecular programs underlying the early evolution of UASCC and establishes genetically defined organoid models as robust tools for mechanistic and therapeutic discovery. These findings provide a foundation for future research aimed at improving early detection and intervention strategies for this aggressive malignancy.

## METHODS

### Mice

All animal procedures were approved by the Institutional Animal Care and Use Committee of the University of Southern California (USC). C57BL/6 mice and athymic nude mice were purchased from the Charles River Laboratories and housed in the USC animal facility.

### Patient samples

HNSCC tumor samples and distal normal tongue tissues were collected from patients undergoing surgery at either Keck Hospital of USC or Yonsei University College of Medicine following Institutional Review Board-approved protocols and with informed consent. Detailed clinical information is available in **Table S1**.

### Cell lines and maintenance

Cultrex HA-R-Spondin-FC 293T cells (3710-001-01) were purchased from Bio-techne and maintained in Dulbecco’s modified Eagle medium (DMEM) with 10% fetal bovine serum (FBS) to generate R-Spondin-1-conditioned medium. HEK293T cells (CRL-1573) were purchased from the American Type Culture Collection and maintained in DMEM with 10% FBS.

### Generation of murine and human organoids

Mouse oral tongue, oropharynx, and esophagi were collected from 6-7-week-old C57BL/6 mice immediately after euthanizing by CO2 inhalation and cervical dislocation. Human surgical specimens were freshly preserved in ice-cold conditioned PBS [PBS containing 2% (v/v) penicillin/streptomycin (P/S) and 1% (v/v) amphotericin B] until further processing within 24 hours. Tissues were rinsed in sterile and ice-cold conditioned PBS [PBS containing 2% (v/v) penicillin/streptomycin and 1% (v/v) amphotericin B] for at least 5 times. Tissues were further dissected to remove the adipose, muscle and necrotic tissues, and the epithelium layer was transferred into tissue processing tubes (RWD) containing 5 ml of digestion buffer [DMEM/Ham’s F-12 containing 2.5% (v/v) FBS, 1% (v/v) P/S, 1 mg/ml collagenase type IX, and 120 μg/ml dispase type II]. Tissue process tubes were then placed in the hushing of the DSC-400 tissue dissociator (RWD), and the 12-minute tissue mincing program and 45-minute digestion program were executed. The suspension was passed through a 70 μm cell strainer, and the flow-through was spun down at 1,000×g for 3 minutes. The sedimented cells were washed 3 times with CPBS and suspended in Cultrex basement membrane extract (BME, Biotech) and dispensed onto 6 - well plates as to 50- or 100-μl droplets. After BME polymerization, the droplets were overlaid with 2 ml of organoid culture medium as described below.

### Organoid culture medium

The culture medium for murine tongue and oropharynx organoids was DMEM/Ham’s F-12 supplemented with 10% (v/v) R-spondin-1 conditioned medium, 10 ng/ml human fibroblast growth factor 10 (rFGF-10), 50 ng/ml mouse recombinant epidermal growth factor (rEGF), 100 ng/ml Noggin, 2.5 mM N-acetylcysteine, 10 mM Nicotinamide, 10 μM Y27632, 1×B-27, 1% (v/v) amphotericin B, 1×P/S, and 1×Primocin. The culture medium for murine esophageal organoids was prepared by supplementing DMEM/Ham’s F-12 50% (v/v) Wnt-3A–conditioned medium, 20% (v/v) R-spondin-1–conditioned medium, 100 ng/ml human rFGF-10, 50 ng/ml mouse rEGF, 100 ng/ml Noggin, 1 mM N-acetylcysteine, 10 mM Nicotinamide, 10 nM gastrin I, 500 nM A-83-01, 10 μM SB202190, 10 μM Y27632, 1×B-27, 1% (v/v) amphotericin B, 1×P/S, and 1×Primocin. The culture medium for human oral organoids was DEME/Ham’s F12 supplemented with 20% R-spondin-1 conditioned medium, 1 μM prostaglandin E2, 10 ng/ml human rFGF10, 50 ng/ml human rEGF, 5 ng/ml human recombinant fibroblast growth factor-basic, 100 ng/ml Noggin, 1 mM N-acetylcysteine, 10 mM Nicotinamide, 500 nM A-83-01, 10 μM SB202190, 10 μM Y27632, 0.3 μM CHIR99021, 1×B-27, 1 μM Forskolin, 1×GlutaMAX, 1×P/S, 1% (v/v) amphotericin B, and 1×Primocin. Detailed reagent information used for organoid culture is listed in **Table S2.**

### Retro- and lenti-virus production

For lentiviral particle generation, 4.5 μg of psPAX2, 1.5 μg of pMD2.G, and 6 μg of the lentiviral construct plasmid (pLV-Bsd-U6-shScramble, pLV-Bsd-U6-shCdkn1b, pLV-Bsd-U6-shCDKN1B, pLentiCRISPRv2-sgCtrl, pLentiCRISPRv2-sgKMT2C, or pLentiCRISPRv2-sgKMT2C) were used. For retroviral particles generation, 6.5 μg of pUMVC (Addgene), 6.5 μg of pCMV-VSV-G (Addgene), and 4.3 μg of retroviral construct plasmid (pBABE-puro or pBabe-puro-HA-PIK3CA-E545K) were used. Plasmids were co-transfected into HEK293T cells at 70% confluency in 10-cm dishes using Lipofectamine 3000 reagent (ThermoFisher Scientific) according to the manufacturer’s protocol. Supernatants containing lentivirus or retrovirus were harvested twice daily for two days. After removing cells by centrifugation, the supernatants were filtered through 0.45 μm syringe filters, aliquoted, and stored at -80°C.

### Genetic engineering and gene editing of organoids

For gene knockout (KO) using Alt-R CRISPR-Cas9 System (Integrated DNA Technologies), organoids were electroporated as previously described (12). Briefly, the Cas9:gRNA RNP complex was prepared as follows: 100 μM gRNA complex was prepared by mixing 200 μM crRNA targeting either murine *Trp53*, murine *Cdkn2a*, human *TP53*, or human *CDKN2A,* 200 μM tracrRNA labeled with ATTO 550. The mixture was heated to 95°C for 5 minutes and then allowed to cool slowly to room temperature; For each electroporation, the RNP complex was prepared by combining 6 μl of the 100 μM gRNA complex, 8.5 μg of Cas9 nuclease, and 10.5 μl of duplex buffer, followed by a 10-minute incubation at room temperature. Organoids were dissociated into clusters of 10-15 cells using TrypLE Express (ThermoFisher Scientific), resuspended in 75 μl of opti-MEM buffer (ThermoFisher Scientific), and mixed with 5 μl of 100 μM electroporation enhancer, and 50 μl of Cas9:gRNA RNP complex. Electroporation was performed using the NEPA21 system (Nepa Gene), with the same parameters previously described (12). The negative control RNP complex was electroporated into organoids for the control group. Two days post-electroporation, organoids were treated with 10 μM Nutlin-3a for 2 weeks to select for *TP53* mutant organoids. Successful introduction of biallelic frameshift mutations was analyzed by the TIDE (68) and further validated by TOPO-TA cloning and Sanger sequencing as previously described (12).

KO of human *KMT2C* or murine *Kmt2c,* as well as knockdown of human *CDKN1B* or murine *Cdkn1b* in organoids were achieved by lentivirus transduction, while *PIK3CA^E545K^* overexpression was achieved through retroviral transduction. For viral transduction, organoids were dissociated into small clusters and suspended in 1 ml of lentiviral or retroviral supernatants containing 8 μg/ml of polybrene. Spinoculation was then performed at 600 × g for 1 hour at 32°C. Following spinoculation, 500 μl of organoid culture medium was added to each well, and the organoids were incubated overnight at 37°C before being reseeded into a new 12-well plate. After two days, organoids were subsequently selected with 2 mg/ml of puromycin for KO of *KMT2C and Kmt2c* and *PIK3CA^E545K^*overexpression, or with 2 μg/ml of blasticidin for knockdown of *CDKN1B* and *Cdkn1b*, for two weeks to enrich for organoids with successful plasmid delivery. To restore human ANXA1 and mouse Anxa1 in organoids, cDNA of human ANXA1 or mouse Anxa1 was cloned into pcDNA3.1+/C-(K)-DYK vector. Organoids were dissociated and resuspended in 100 μl of electroporation buffer containing 4 μM electroporation enhancer and 10 μg of vector. Electroporation was performed into the organoids as previously described (69). Two days following electroporation, organoids were selected with 400 μg/ml G418 for one week. All reagents and sequences of crRNA, sgRNA, and shRNA used in this section are listed in **Table S2**.

### Organoid viability assay (WST-1 assay)

To quantify metabolically-active viable cells, organoids were seeded and cultured in 96-well plates. At specified time points, a mixture of cell proliferation reagent WST-1 (Cell Biolabs) and fresh organoid culture medium, prepared at a 1:10 ratio, was added to each well (100 µl per well). The organoids were incubated with the mixture at 37℃ for 90 minutes. Post-incubation, only the medium was transferred to a new 96-well plate, and absorbance was measured at 450 nm using a microplate absorbance reader (Accuris).

### Organoid invasion assay

Organoids were dissociated into single cells and resuspended in 60% Cultrex Spheroid Invasion Extracellular Matrix (Bio-Techne) at a density of 2×10^4^ cells/ml. A 50 μl aliquot of the cell suspension was seeded into each well of ultra-low binding 48-well plates. After a 10-minute incubation at 37°C to facilitate matrix polymerization, 500 μl of pre-warmed organoid culture medium was added to each well. The invasiveness of organoids grown in the invasion extracellular matrix was determined by the formation and enrichment of spindle-like protrusions. Brightfield images of the organoids were captured using a Keyence BZX810 microscope on day 10 post-culture.

### RNA extraction and quantification of PCR

Total RNA was extracted from organoids using the RNeasy Mini Kit (Qiagen) and synthesized into cDNA using the LunaScript™ RT SuperMix (New England Biolabs) following manufacturers’ protocol. Real-time qPCR was performed using PowerUp™ SYBR™ Green Master Mix (ThermoFisher Scientific) on CFX96 qPCR System (BioRad). Relative gene expression levels were calculated with the ΔΔCT method. Primers used for real-time qPCR are listed in **Table S2**.

### Allografting of murine organoids

For mouse allograft experiments, 5-6-week-old male athymic nude mice were randomized into control/experimental groups. Three days before transplantation, organoids were passaged following standard protocols. On the day of transplantation, cells were resuspended in cold 50% BME/PBS at a density of 2 × 10^7^ cells per ml. Mice were subcutaneously injected with organoids into both sides of the axillary region (2 × 10^6^ cells per injection). Allograft sizes were monitored by measuring the length (L) and width (W) with a caliper, and the volume (V) was calculated using the formula V = 0.5236 × (L × W^2^). Mice were sacrificed 81 and 84 days post-injection unless otherwise specified, and allografts were isolated for subsequent analysis.

### Histology analysis, H&E, and IF staining

For cryosections, tissues or organoids were fixed in 4% paraformaldehyde in PBS at 4°C overnight, washed twice with PBS at room temperature, dehydrated in 30% sucrose in PBS, and embedded in Tissue-Tek O.C.T. Compound (Sakura). For preparation of paraffin-embedded samples, fixed tissues were processed using Epredia STP 120 Spin Tissue Processor (ThermoFisher Scientific) and manually embedded following standard protocols. O.C.T- or paraffin-embedded samples were sectioned into 6-μm slices. H&E staining was performed using the H&E Staining Kit (Abcam) according to manufacturer’s instructions. For IF staining, the following primary antibodies were used: chicken anti-Krt5 (1:1000, Biolegend), rabbit anti-Krt13 (1:1000, Proteintech), mouse anti-Ki67 (1:1000, Cell Signaling Technology), rabbit anti-Ki67 (1:1700, Abcam), chicken anti-Krt14 (1:500, Biolegend), rabbit anti-Klf4 (1:500, Proteintech), rabbit anti-Anxa1 (1:1000, Abcam) antibodies. Secondary labeling was performed with Alexa Fluor 488-, 568-, or 647-conjugated anti-mouse, or anti-rabbit antibodies (1:1000, ThermoFisher Scientific), or FITC-conjugated anti-chicken antibodies (Biolegend). Fluoroshield with DAPI (ThermoFisher Scientific) were used for nuclei counterstained and mounting. Images were captured using a BZ-X810 digital microscope (Keyence). Details of antibodies and assay kits can be found in **Table S2**.

### SA-β-gal staining

To detect cellular senescence in organoids, the β-galactosidase Staining Kit (Cell Signaling Technology) was used according to manufacturer’s instructions. Organoid-BME domes were washed twice with PBS, fixed for 20 minutes with 4% paraformaldehyde in PBS, and then washed twice with PBS. O.C.T slides were directly washed with PBS. Organoids or slides were then incubated for 18 hours at 37°C with a Staining Solution containing 1 mg/ml X-Gal in DMSO (pH 6.0) and imaged using a BZ-X810 digital microscope (Keyence).

### Protein analysis

Western blotting was performed according to standard protocols. Briefly, organoid pellets were lysed in RIPA Lysis and Extraction Buffer containing Halt Protease Inhibitor Cocktail and Halt Phosphatase Inhibitor Cocktail (ThermoFisher Scientific). Protein concentrations were determined using the BCA Protein Assay Kit (ThermoFisher Scientific). Protein lysates were separated on 12% SurePAGE gels (Genscript) and transferred onto 0.2-μm PVDF membranes (Millipore). After blocking with 5% nonfat dry milk or 5% BSA in 1×TBS buffer (Sigma) with 0.05% Tween-20, the membranes were incubated with the following primary antibodies: rabbit anti-Anxa1 (1:3000), mouse anti-actin (1:3000), rabbit anti-p27 (1:1000, Invitrogen), rabbit anti-SMAD2/3 (1:1000, Cell Signaling Technology), and rabbit anti-pSMAD2/3 (1:1000, Cell Signaling Technology). Secreted Anxa1 and Ecm1 protein levels were determined by ELISA kits (Abcam). Organoids were cultured in 12-well plates for 48 hours. Collected media were centrifuged at 500×g to remove cells and debris. The supernatants were aliquoted and stored at -80°C. The ELISA was performed following the manufacturer’s protocol using 200 μl of sample per well.

### Chromatin immunoprecipitation

ChIP experiments were conducted using the EZ-Magna ChIP A/G Kit (Merck Millipore, 17-10086) following the manufacturer’s protocol. Organoid derived cells were cross-linked with 1% formaldehyde and lysed using the cell lysis buffer containing 1×Protease Inhibitor Cocktail II (PIC II). Nuclei were isolated with the nuclear lysis buffer supplemented with 1×PIC II. The chromatin extract was sonicated and sheared to lengths between 200 and 800 bp (for 8 minutes in total, 30% amplitude, pulse on for 10 s, and pulse off for 20 s). Sheared, cross-linked chromatin was immunoprecipitated with antibodies incubated with magnetic protein A/G beads. Antibodies used included anti-pSMAD3 (6 μg per ChIP reaction; Invitrogen) and normal mouse IgG (1 μg per reaction). Purified DNA was subjected to qPCR, with primers listed in **Table S2.**

### Whole exome sequencing (WES) and data analysis

Genomic DNA was extracted from patient tissues and matched peripheral blood samples using the DNeasy Blood & Tissue Kit (Qiagen). DNA libraries were prepared with the Agilent SureSelect Human All Exon V6 Kit (Agilent) and sequenced on the Illumina NovaSeq 6000 platform with paired-end 150 bp reads. Quality control of raw reads was performed using FastQC, followed by the removal of low-quality reads and adapters. Clean paired-end reads were aligned to the human reference genome (GRCh38) using the Burrows-Wheeler Aligner (BWA). Single-nucleotide variants (SNVs) and small insertions and deletions (InDels) were identified using the Genome Analysis Toolkit (GATK)(70). Variants were annotated with ANNOVAR(71).

### Single-Cell RNA-Sequencing and data processing

Mouse organoids were dissociated into single cells and resuspended in organoid culture medium for the library preparation. The Chromium Next GEM Single-Cell 3ʹ Reagent Kit (10× Genomics) was used according to the manufacturer’s protocols and the procedure was conducted by the Fulgent Genetics. Raw reads were aligned to the mouse GRCm38/mm10 reference genome and unique molecular identifiers (UMIs) were estimated using the Cell Ranger v.7.0.1 (10× Genomics). The Seurat (v4.3.0.1) R package (72) was used to filter the raw data of the gene expression matrix. Cells with UMIs ≥500, genes ≥200 and ≤8000, log10 (Genes Per UMI) > 0.80, and mitochondrial gene content < 10%, and genes expressed in more than 1/1000 cells were retained. Doublets were removed using the R packages DoubletFinder (v2.0.3) (72,73). After sequencing quality control and standardization, there were a total of 6979 cells from 3 mouse organoid lines, with an average of 18,391 genes being measured in each mouse organoid line. The batch effects of the samples were corrected using the ‘RunHarmony’ function in the “harmony” (v0.1.1) R package (72–74). The cells were clustered using UMAP with the ‘RunUMAP’ function (dims = 1:30), and the cluster-specific marker genes were identified using the Seurat function ‘‘FindAllMarkers’’ with the default arguments.

### Non-negative matrix factorization (NMF) analysis and identification of gene programs

NMF was employed to analyze each mouse organoid line individually, to identify gene modules or programs. All negative values in the centered expression matrix were set to zero as previously described (31). NMF was performed with a range of factors (K) from 6 to 9, resulting in 30 programs per mouse organoid line. Each program was characterized by the top 50 genes based on NMF coefficients. To identify robust programs across different K values and among the three mouse organoid lines, we applied three criteria: 1) A program had to show at least 70% overlap (35 of 50 genes) with a program of a different K value within the same organoid line; 2) A program had to share at least 20% overlap (10 of 50 genes) with a program in any other organoid line; 3) The programs were ranked based on their similarity with gene programs from other organoid lines, and selection was performed in decreasing order. Once a program was chosen, any other program within the same organoid line that had >20% overlap with the selected gene program was excluded to reduce redundancy. This process resulted in 19 robust programs, which were then subjected to hierarchical clustering using the unweighted pair group method with arithmetic mean based on Jaccard similarity, and finally 7 gene programs (GPs) were manually defined. For each GP, genes present in at least 40% of its constituent GPs were defined as program signature genes. The gene lists for each GP are provided in **Table S3**. To understand the identified GPs, we compared them against 41 curated GPs (termed meta-programs or MPs by the original study) established by the same NMF approach from scRNA-seq delineation of pan-cancer tumor samples (29). We calculated the average similarity (Jaccard index) and single-cell score correlation between each MP and our GPs. Statistical significance was assessed by permuting our programs 100 times, with a 99.9% confidence threshold.

### Single-Cell Gene Program Scoring

For each of the 7 consensus GPs, 1,000 background gene sets with comparable expression levels were randomly generated as previously described (75). For each single cell, we measured the average centered expression of the GP gene sets and the 1,000 randomly generated gene sets. The p-value was defined as the proportion of randomly generated gene sets that was expressed at higher levels than the GP gene sets. The GP score was calculated as -log10(p) and linearly rescaled to a range of [0,1]. To directly compare our mouse organoid single-cell gene program scores with HNSCC patient tumors (25) and patient tumor-derived organoid (PDO) data (10), we selected samples harboring the EMT and EpiSen programs and integrated our mouse organoids into joint datasets(25). Expression levels (log2(TPM+1)) of each dataset were mean-centered per gene before combining the cells. Principal component analysis (PCA) was performed on the joint datasets using the top 4,500 genes with the highest expression levels. For visualization, we selected the top two principal components (PCs) combinations that showed a strong correlation (r > 0.35 or r < -0.35) with the single-cell scores corresponding to the GPs of interest.

### Identification of Representative Genes for GP7

To define a group of representative genes for GP7, we first focused on the 23 genes identified as overlapping with the SASP database (**Fig. 4C**). Among these, Anxa1, Ecm1, Il1rn, and Cxcl5 emerged as key candidates, each exceeding a log₂ fold-change threshold (>2) in at least one senescence condition and exhibiting a detection frequency of 1 in the mouse scRNA-seq data. We next checked the overlap genes between GP7 and our epithelial senescence GP5 or HNSCC patient tumors EpiSen program and revealed two additional overlapped genes Cxcl1 and Cldn4 (**Fig. S16**). These genes were identified as representative genes of epithelial senescence and secretion for further study.

### Drug screening at single-organoid resolution via bioprinting coupled with machine learning-assisted imaging analysis

High throughput drug screening for organoids was conducted as previously described with minor modifications. Briefly, we used a volumetric temperature-controlled printer (Corning) for bioprinting. A bespoke python script was used to integrate coordinates of well centers, print patterns (rings), ring size (5.9 mm in diameter) and volume (10 μl) into G-code files to achieve the desired single-layer geometry. On day 0, mouse organoids were passaged and dissociated into single cells. We prepared a bioink composed of single-cell suspension in a 3:4 mixture of organoid culture medium and Matrigel (Corning) on ice. After briefly vortexing, the mixture was transferred into a 3 ml syringe with a 20-gauge needle and loaded into the bioprinter. We used glass-bottom plates pre-treated with oxygen plasma using PE-25 Plasma Etcher (Plasma Etch) for 90 seconds for printing. As we previously reported, we took advantage of 3D-printed masks to achieve optimal print location and thickness (55). After printing, constructs were incubated at 37°C for at least 30 minutes to solidify the matrix, followed by the addition of 100 μl of organoid culture medium. On day 4 and 5, we performed drug treatments by performing full media exchanges. All drugs were dissolved in DMSO, which was used as control. Lastly, 24h after the last drug treatment, we released the organoids from the matrix using 50 µl of a 5 mg/ml dispase solution (Life Technologies) and performed an end-point ATP-release luminescent assay using CellTiter-Glo 3D Luminescent Cell Viability Reagent (Promega) following manufacturer’s instructions. Luminescence was measured with a SpectraMax iD3 plate reader (Molecular Devices), and viability was determined by normalizing the signal from each well to the average signal of the negative control wells (DMSO). Z scores were calculated using the formula: (average ATP assay value of a drug treatment − average ATP value of the DMSO control) / standard deviation of the DMSO control.

For the entire duration of the experiment, we acquired daily brightfield images of organoids using a Celigo imager, with 3 z-planes imaged in each well at a focal height difference of 75 µm. The organoid size was measured and normalized from these projected areas, and segmented using machine learning-based models as previously described (52). The normalized organoid growth rates were determined by comparing the average organoid size on day 6 (prior to the ATP assay) to the average size on day 4 (prior to the initial drug treatment), with both values normalized to the control (DMSO vehicle).

### Drug treatment of human organoids

To validate and test a range of candidate drugs identified from high-throughput screenings on mouse organoids, human organoids were cultured at a density of 1,000 cells per 15 μl BME in 96-well plates, followed by the addition of 100 μl of organoid culture medium. After a three-day recovery period for organoid formation, brightfield images were captured using a BZ-X810 digital microscope. The medium was then completely removed, and the organoids were treated with serial dilutions of the drugs, ranging from 0.05 μM to 1280 μM, with DMSO-treated organoids serving as the control group. Four days post-treatment, additional brightfield images were taken, followed by the WST-1 assay as described above. The combination index for Mitomycin C and Onalespib was calculated using the Chou-Talalay method (76).

### Statistical analysis

All experiments reported in this study were repeated at least three independent times, unless otherwise specified in the figure legends or Methods. All sample numbers (n) of biological replicates and technical replicates can be found in the figure legends. The data are presented as means ± SD and statistical analysis was assessed using GraphPad Prism 10.2 unless otherwise stated. *P* < 0.05 was considered statistically significant. Further statistical details are indicated in each figure legend.

## Data availability

The raw and processed data generated in this study have been deposited in the database under accession code GSE286449. Single-cell datasets and bulk RNA-seq data used in the study are obtained from: GSE234933, GSE181919, GSE182227, GSE198315, HRA004648, as well as TCGA-HNSCC (https://portal.gdc.cancer.gov/). Source data are provided with this paper.

## Authors’ Contributions

**H. Z.:** Conceptualization, formal analysis, validation, investigation, visualization, methodology, writing–original draft, writing–review and editing, data curation. **Y.-M. P.:** Resources, software, visualization. **Y. Z:** formal analysis, software, visualization, methodology. **Q. M.:** formal analysis, software, visualization, methodology. **C. C.:** Investigation. **B. H.:** Investigation. **T. Z.:** Investigation, visualization. **L. L.:** Investigation, visualization, Methodology. **S. W.:** Investigation. **Y. P.:** Investigation. **A.V. M.:** Resources. **U.K. S.:** Supervision, resources, funding acquisition. **P. S.:** Validation. **A. S.:** Supervision, resources, data curation, funding acquisition, project administration, writing–review and editing. **D.-C. L.:** Conceptualization, supervision, resources, data curation, funding acquisition, project administration, writing–original draft, writing–review and editing.

## Acknowledgments

We extend our gratitude to the patients who participated in this study. Figure 1B and Figure 2B were created with BioRender software (https://biorender.com/).

## SUPPLEMENTS

**Figure S1.**
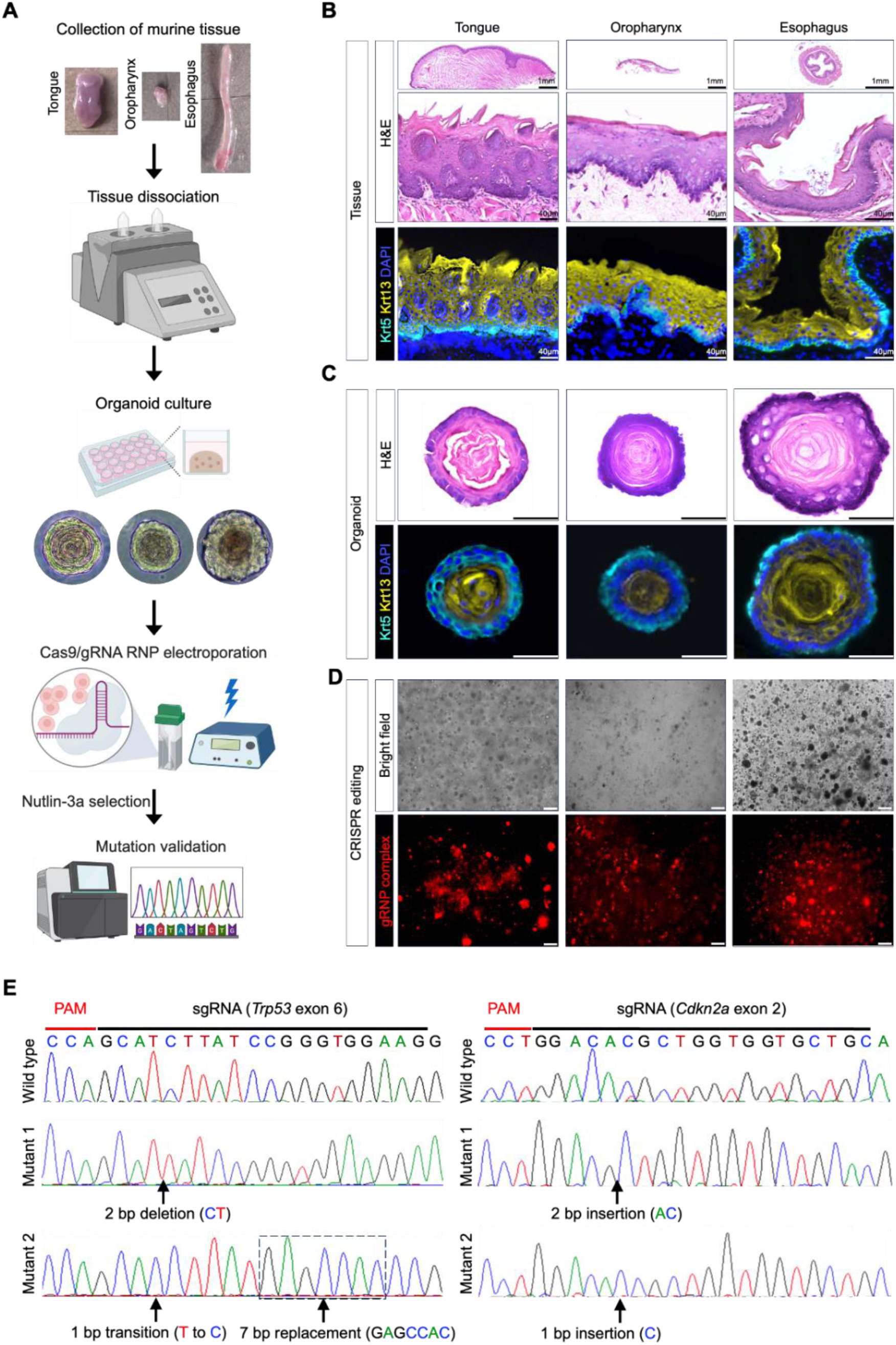
Establishment of CRISPR-Cas9 knockout organoid lines from murine upper aerodigestive tissues. **(A)** Workflow for generating and validating CRISPR-Cas9 edited organoids from mouse tissue. (**B)** Representative H&E and IF images showing staining for the basal cell marker Krt5 (aqua) and the squamous differentiation marker Krt13 (yellow) in mouse tissues. **(C)** Representative H&E and IF images showing staining for Krt5 (aqua) and Krt13 (yellow) in mouse organoids. **(D)** Organoids transfected with Cas9 nuclease and either a negative control gRNA or *Trp53*/*Cdkn2a*-targeted gRNA complex, visualized by red fluorescence. Scale bar, 100 μm. **(E)** Sanger sequencing showing representative mutations at targeted sites, with PAM sequences underlined in red in wild-type sequences. Scale bar, 100 μm.

**Figure S2.**
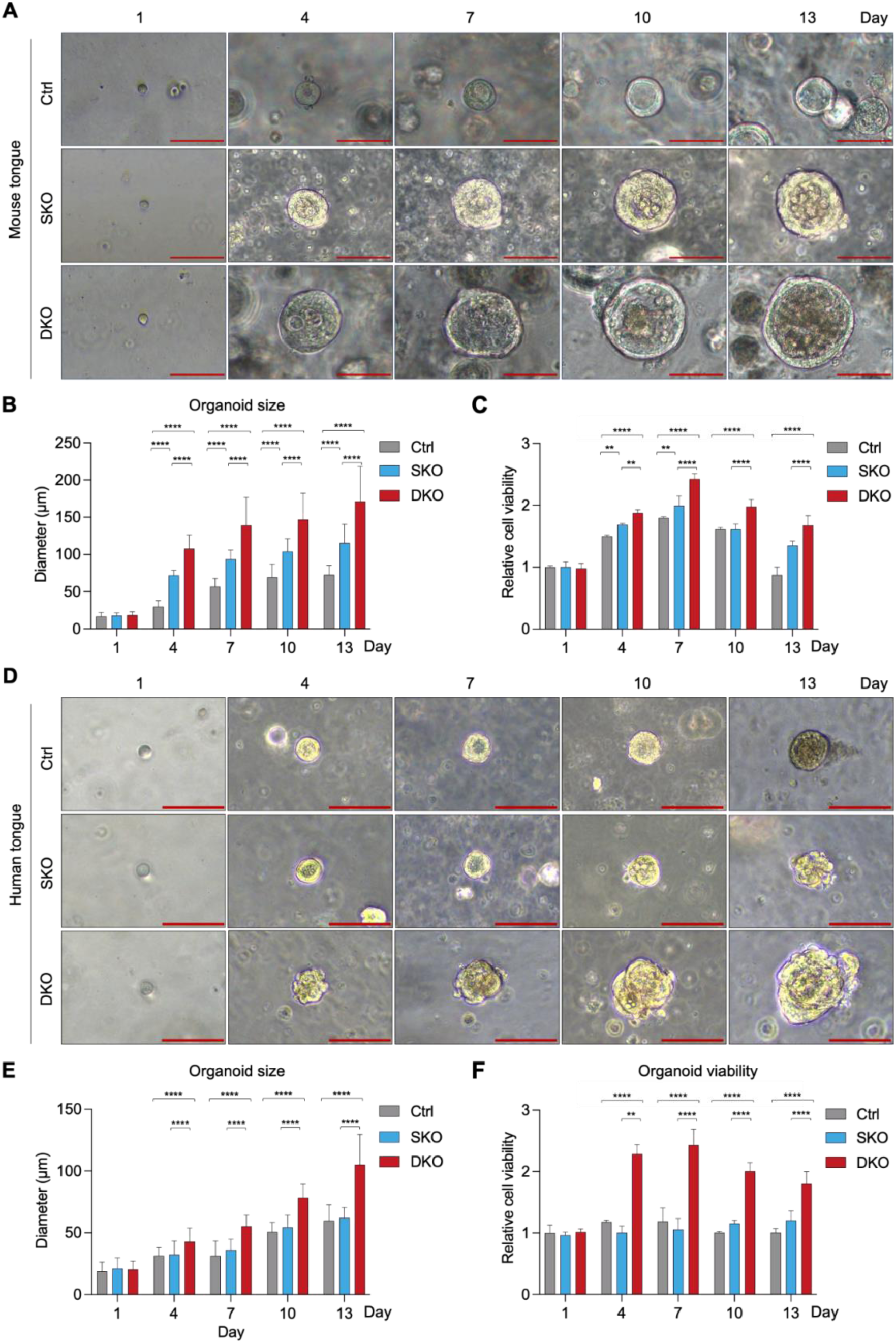
Growth kinetics of genetically-engineered mouse and human tongue organoid lines. (A-C) Mouse tongue Ctrl, SKO and DKO organoids were analyzed for growth properties at the indicated time points: **(A)** Representative phase-contrast photomicrographs of mouse organoid lines. **(B)** Average size of mouse organoids measured at each time point (n = 50 per group). **(C)** Viability of mouse organoids assessed by the WST-1 assay (n = 6 per group). **(D-F)** Human tongue Ctrl, SKO and DKO organoids were analyzed for growth properties at the indicated time points: **(D)** Representative phase-contrast photomicrographs of human organoid lines. **(E)** Average size of human organoids measured at each time point (n = 50 per group). **(F)** Viability of human organoids assessed by the WST-1 assay (n = 6 per group). Scale bar, 100 μm. \*\**P* < 0.01; \*\*\*\**P* < 0.0001.

**Figure S3.**
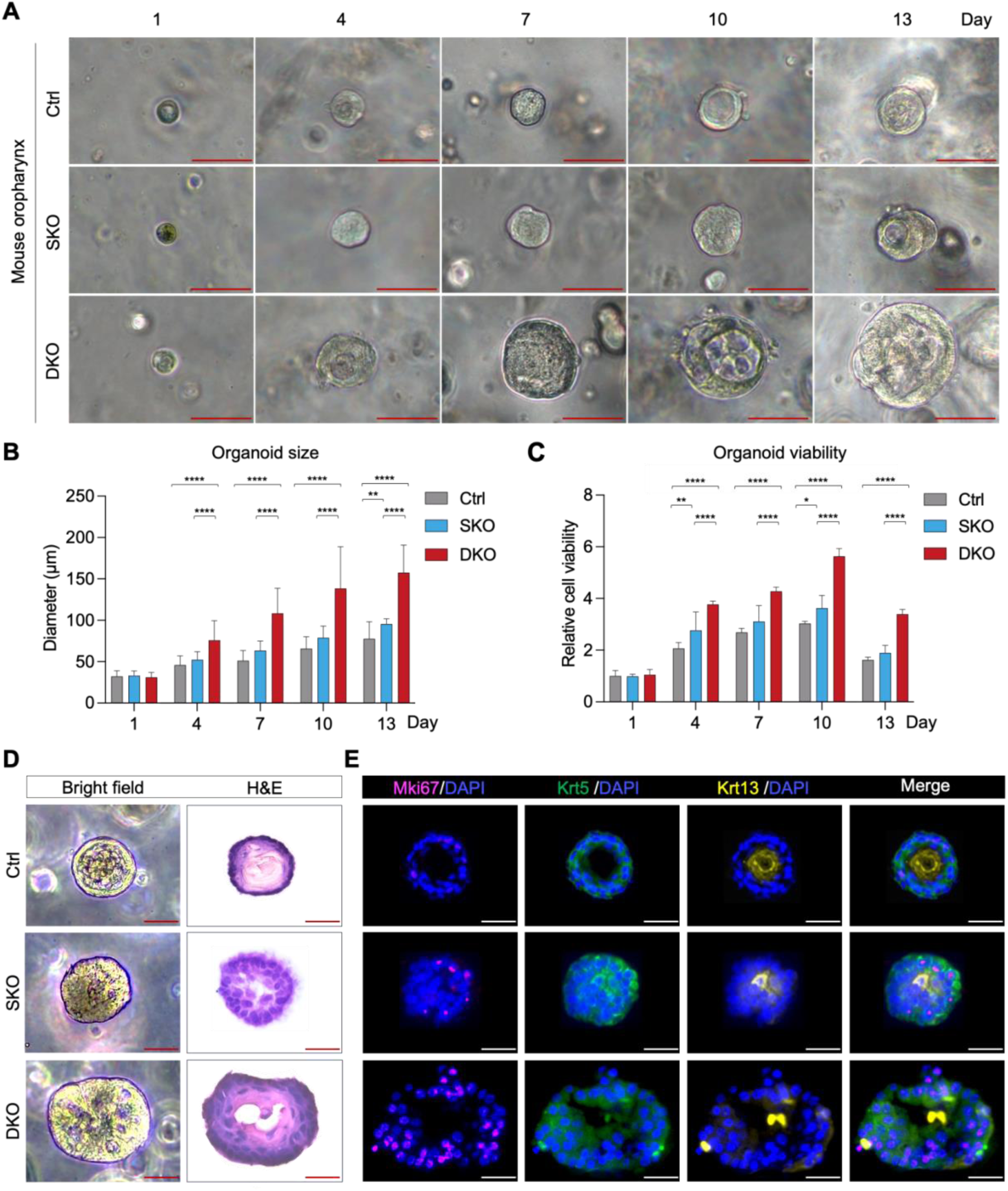
Growth kinetics of genetically-engineered mouse oropharynx organoid lines. **(A)** Representative phase-contrast photomicrographs of mouse oropharynx organoid lines. **(B)** Average size of mouse oropharynx organoid lines measured at each time point (n = 50 per group). (C) Viability of mouse oropharynx organoid lines assessed by the WST-1 assay (n = 6 per group). **(D)** Representative bright-field, H&E, and (**E**) IF images showing staining for the proliferation marker Mki67 (magenta), basal cell marker Krt5 (green), and squamous differentiation marker Krt13 (yellow) in 3-week-old mouse oropharynx Ctrl, SKO, and DKO organoids. Scale bar, 100 μm. \**P* < 0.05; \*\**P* < 0.01; \*\*\*\**P* < 0.0001.

**Figure S4.**
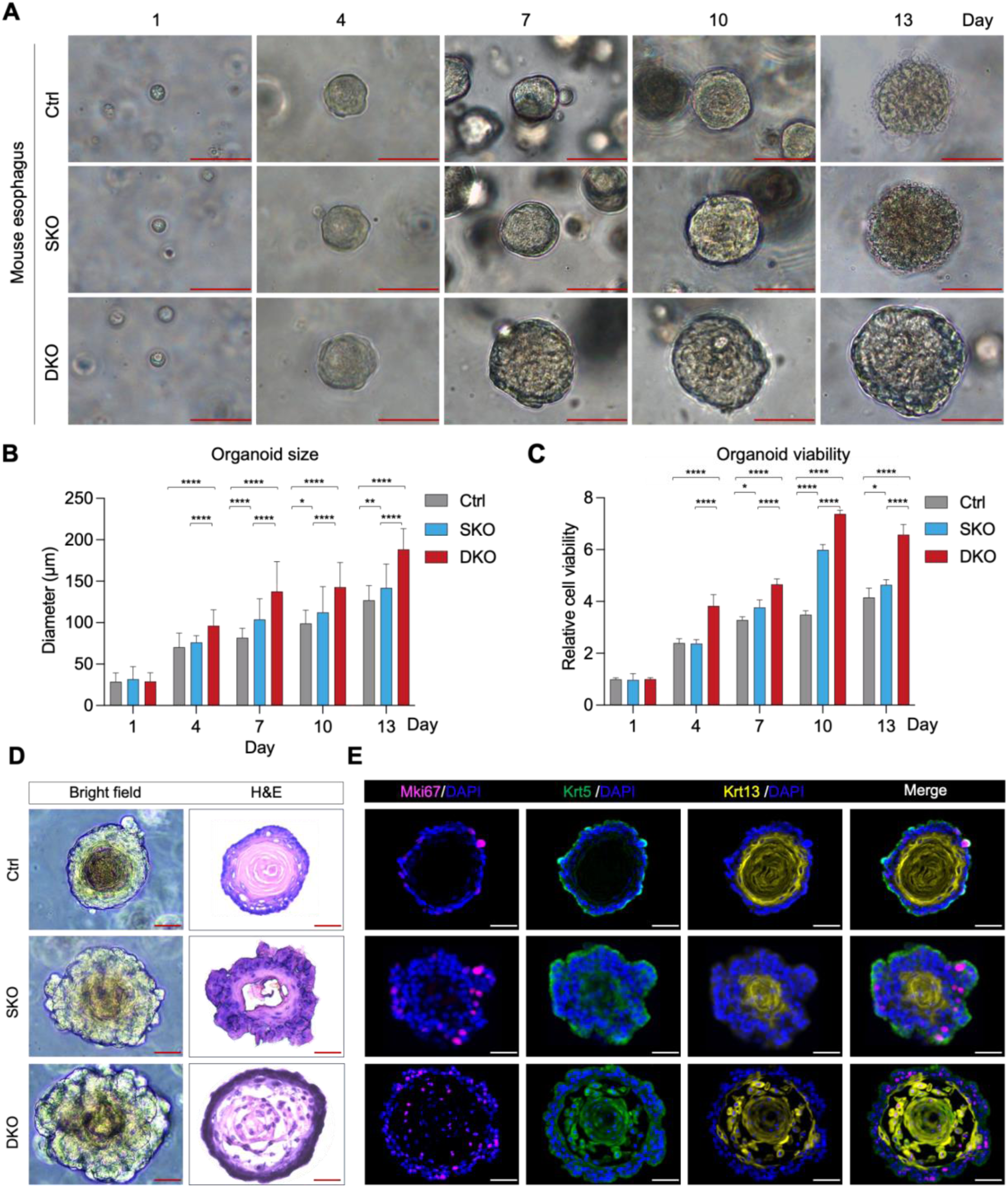
Growth kinetics of genetically-engineered mouse esophageal organoid lines. **(A)** Representative phase-contrast photomicrographs of mouse esophageal organoid lines. **(B)** Average size of mouse esophageal organoid lines measured at the indicated time points (n = 50 per group). **(C)** Viability of mouse oropharynx organoid lines assessed by the WST-1 assay (n = 6 per group). **(D)** Representative bright-field, H&E, and **(E)** IF images showing staining for the proliferation marker Mki67 (magenta), basal cell marker Krt5 (green), and squamous differentiation marker Krt13 (yellow) in 3-week-old mouse esophageal Ctrl, SKO, and DKO organoids. Scale bar, 100 μm. \**P* < 0.05; \*\**P* < 0.01; \*\*\*\**P* < 0.0001.

**Figure S5.**
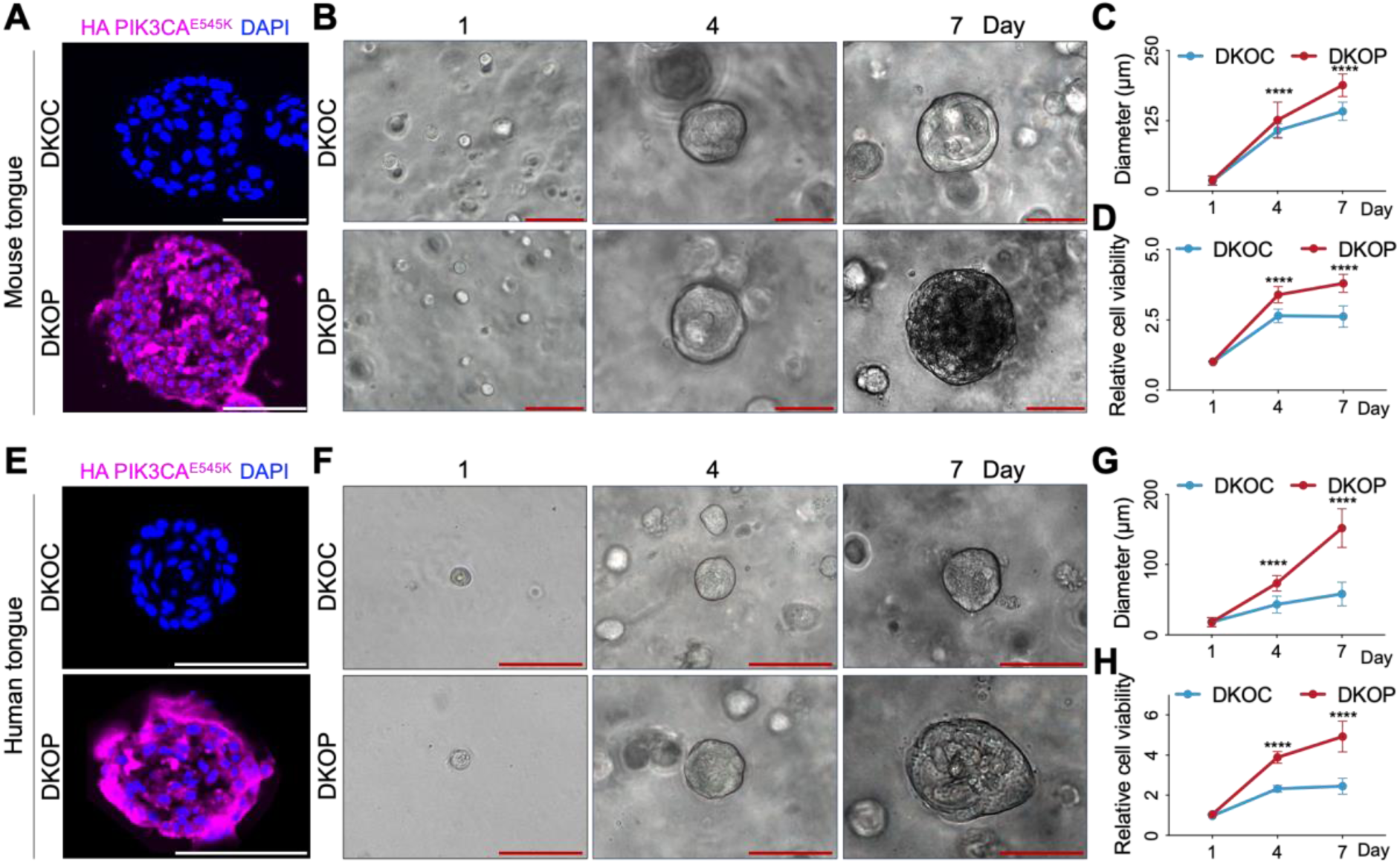
Analysis of PIK3CA^E545K^ expression and growth properties in mouse and human tongue DKOC and DKOP organoids. (A-D) Mouse tongue DKOC and DKOP organoids: **(A)** IF staining images showing HA-PIK3CA^E545K^ (magenta) expression in mouse organoid lines. **(B)** Representative phase-contrast photomicrographs of mouse organoid lines. **(C)** Average size of mouse organoid lines measured at the indicated time points (n = 50 per group). **(D)** Viability of mouse organoid lines assessed by the WST-1 assay (n = 6 per group). **(E-H)** Human tongue DKOC and DKOP organoids: **(E)** IF staining images showing HA-PIK3CA^E545K^ (magenta) expression in human organoid lines. **(F)** Representative phase-contrast photomicrographs of human organoid lines. **(G)** Average size of human organoid lines measured at the indicated time points (n = 50 per group). **(H)** Viability of human organoid lines assessed by the WST-1 assay (n = 6 per group). Scale bar, 100 μm. \*\*\*\**P* < 0.0001.

**Figure S6.**
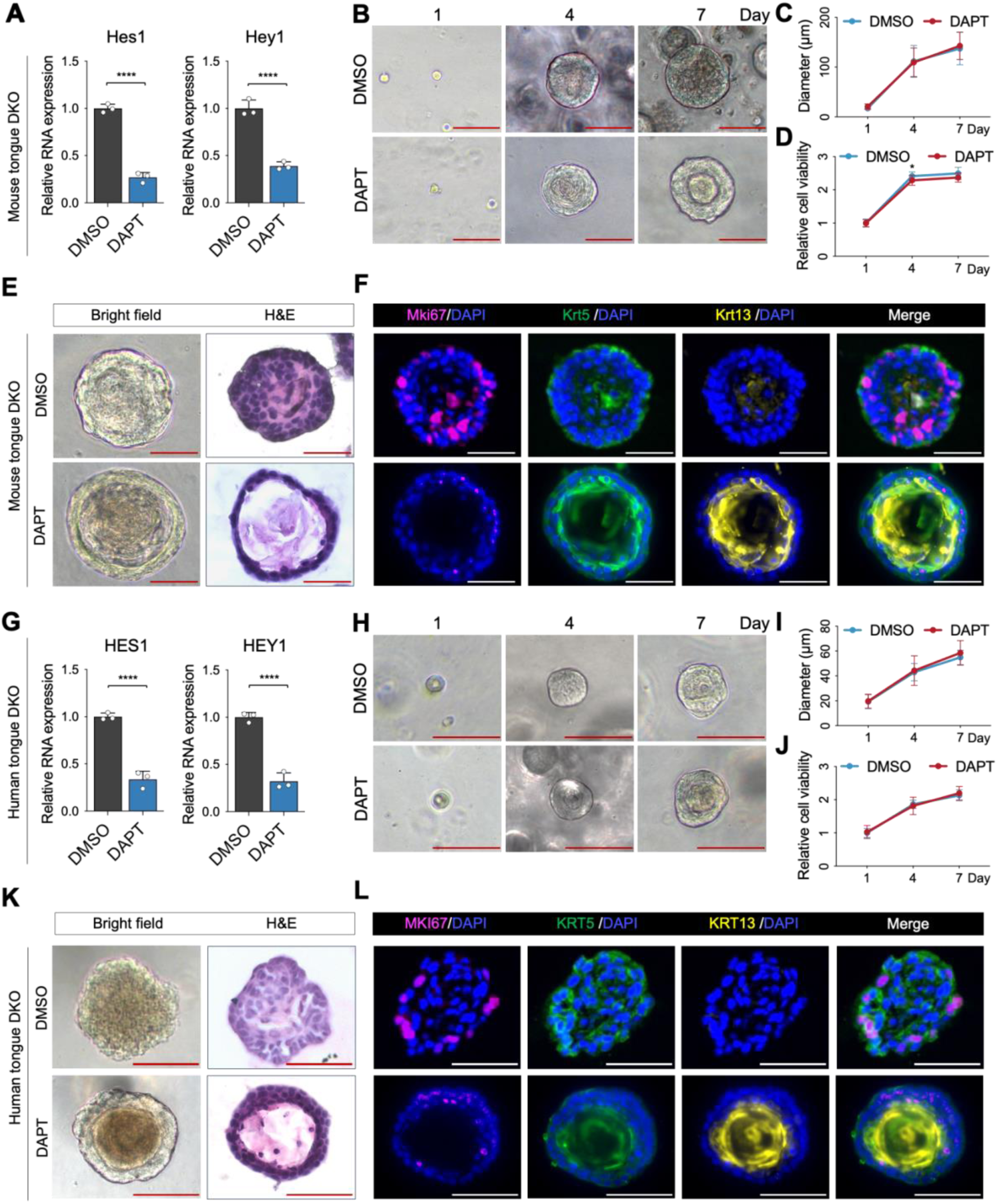
Analysis of NOTCH inhibition and growth properties in human tongue DKOC and DKOP organoids. (A-F) Mouse tongue DKO organoids treated with DMSO or the NOTCH inhibitor DAPT: **(A)** Relative mRNA expression levels of key Notch signaling-related genes. **(B)** Representative phase-contrast photomicrographs of mouse organoid lines. **(C)** Average size of mouse organoids measured at the indicated time points (n = 50 per group). **(D)** Viability of mouse organoids assessed using the WST-1 assay (n = 15 per group). **(E)** Representative bright-field and H&E images, and **(F)** IF images showing staining for the proliferation marker Mki67 (magenta), basal cell marker Krt5 (green), and squamous differentiation marker Krt13 (yellow) in 3-week-old mouse organoids. **(G-L)** Human tongue DKO organoids treated with DMSO or DAPT: **(G)** Relative mRNA expression levels of key NOTCH signaling-related genes. **(H)** Representative phase-contrast photomicrographs of human organoid lines. **(I)** Average size of human organoids measured at the indicated time points (n = 50 per group). **(J)** Viability of human organoids assessed using the WST-1 assay (n = 15 per group). **(K)** Representative bright-field and H&E images, and **(L)** IF images showing staining for the proliferation marker MKI67 (magenta), basal cell marker KRT5 (green), and squamous differentiation marker KRT13 (yellow) in 3-week-old human organoids. Scale bar, 100 μm. \*\*\*\**P* < 0.0001.

**Figure S7.**
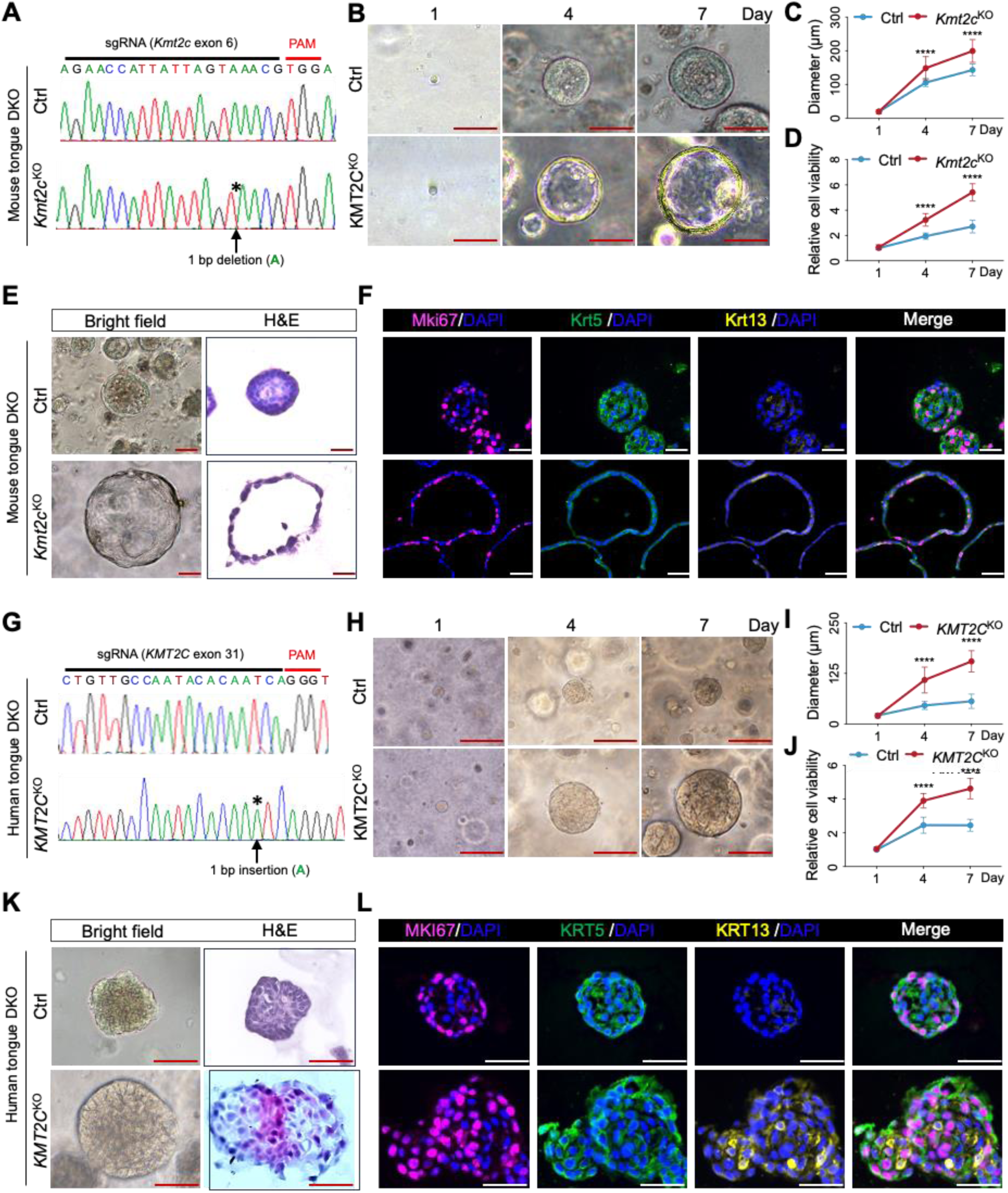
Analysis of targeted *KMT2C* knockout and growth properties in mouse and human organoids. (A-F) Mouse tongue DKO organoids with Ctrl (non-targeting sgRNA) or *Kmt2c* knockout: **(A)** Sanger sequencing showing representative mutations at the targeted Kmt2c sites. **(B)** Representative phase-contrast photomicrographs of mouse organoid lines. **(C)** Average size of mouse organoids measured at the indicated time points (n = 50 per group). **(D)** Viability of mouse organoids assessed using the WST-1 assay (n = 6 per group). **(E)** Representative bright-field and H&E, and **(F)** IF images showing staining for the proliferation marker Mki67 (magenta), basal cell marker Krt5 (green), and squamous differentiation marker Krt13 (yellow) in 3-week-old mouse organoids. **(G-L)** Human tongue DKO organoids with Ctrl or *KMT2C* knockout: **(G)** Sanger sequencing showing representative mutations at the targeted *KMT2C* sites. **(H)** Representative phase-contrast photomicrographs of human organoid lines. **(I)** Average size of human organoids measured at the indicated time points (n = 50 per group). **(J)** Viability of human organoids assessed using the WST-1 assay (n = 6 per group). **(K)** Representative bright-field and H&E images of 3-week-old human organoids. **(L)** IF images showing staining for MKI67 (magenta), KRT5 (green), and KRT13 (yellow) in 3-week-old human organoids. Scale bar, 100 μm. \*\*\*\**P* < 0.0001.

**Figure S8.**
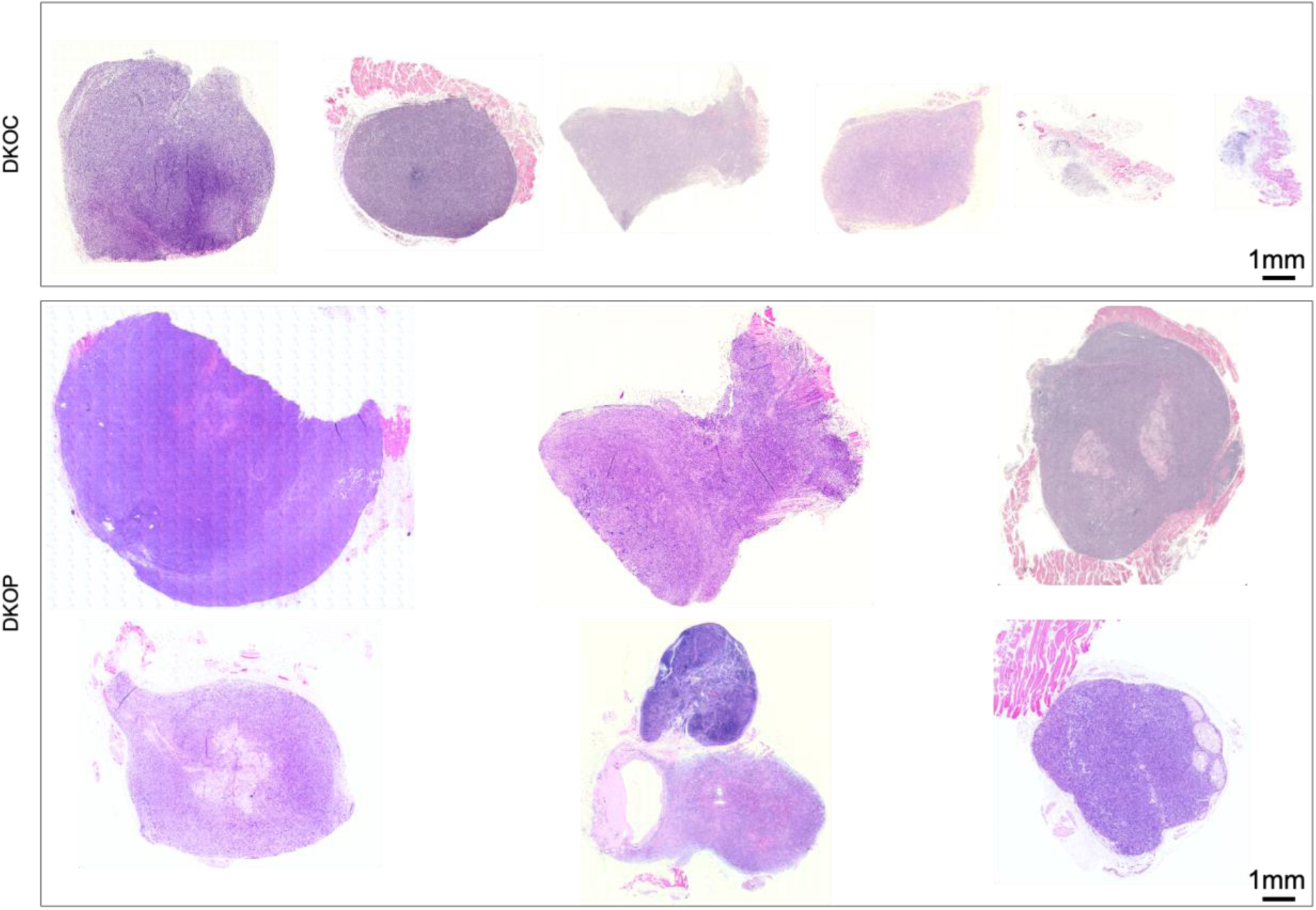
H&E staining of tumors formed by mouse tongue DKOC and DKOP organoids in vivo.

**Figure S9.**
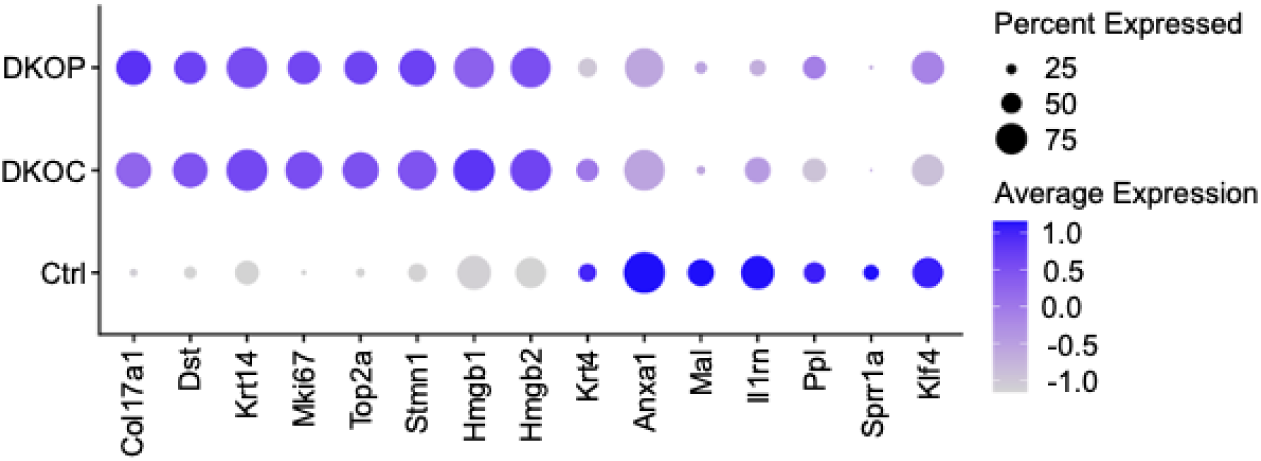
Dot plot showing the expression of selected marker genes in mouse tongue organoid lines.

**Figure S10.**
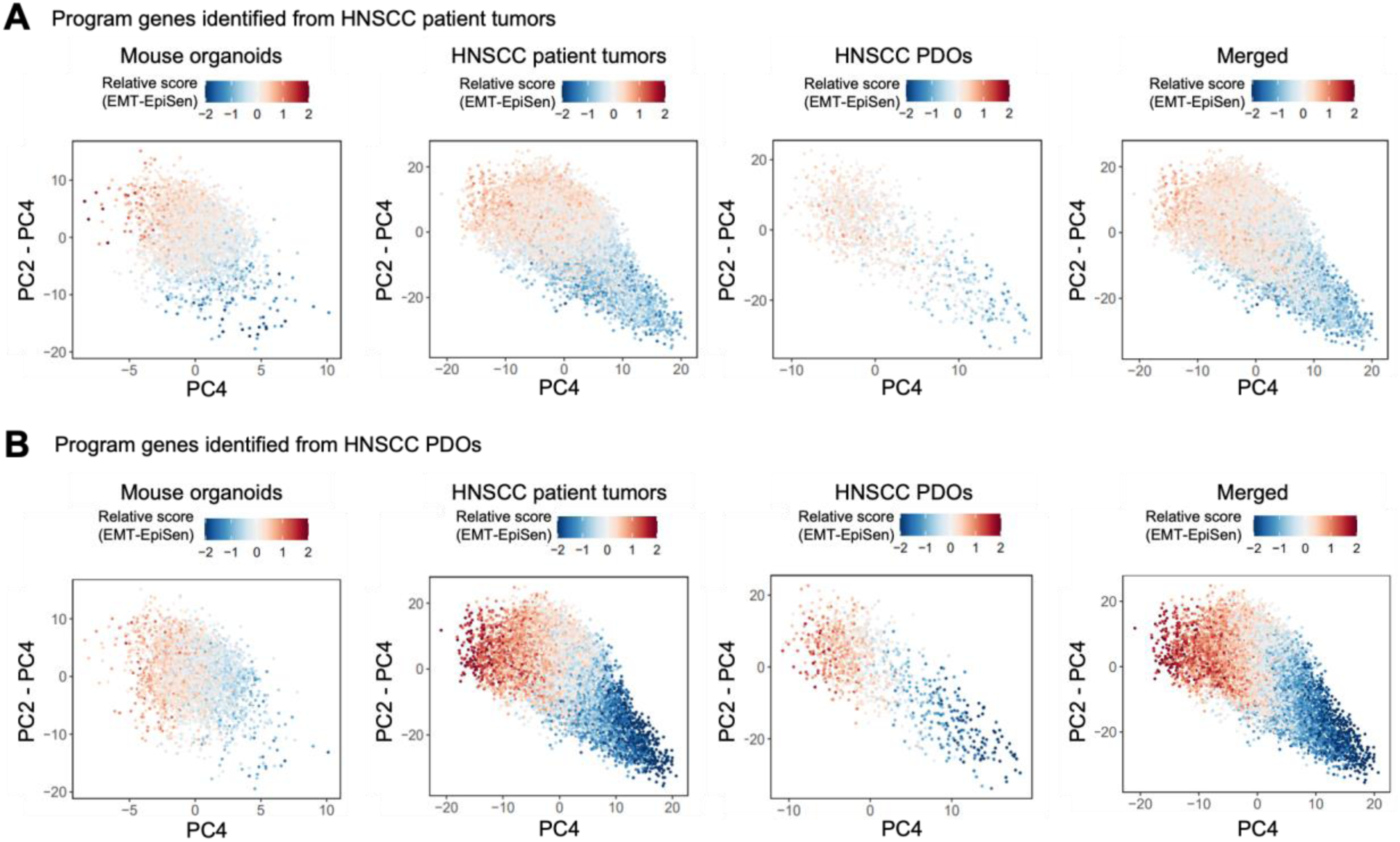
PCA plots of indicated single-cell samples. Cells from mouse organoids, HNSCC patient tumors, and HNSCC PDOs are colored by relative scores for EMT and EpiSen genesets identified in **(A)** HNSCC patient tumors and **(B)** HNSCC PDOs.

**Figure S11.**
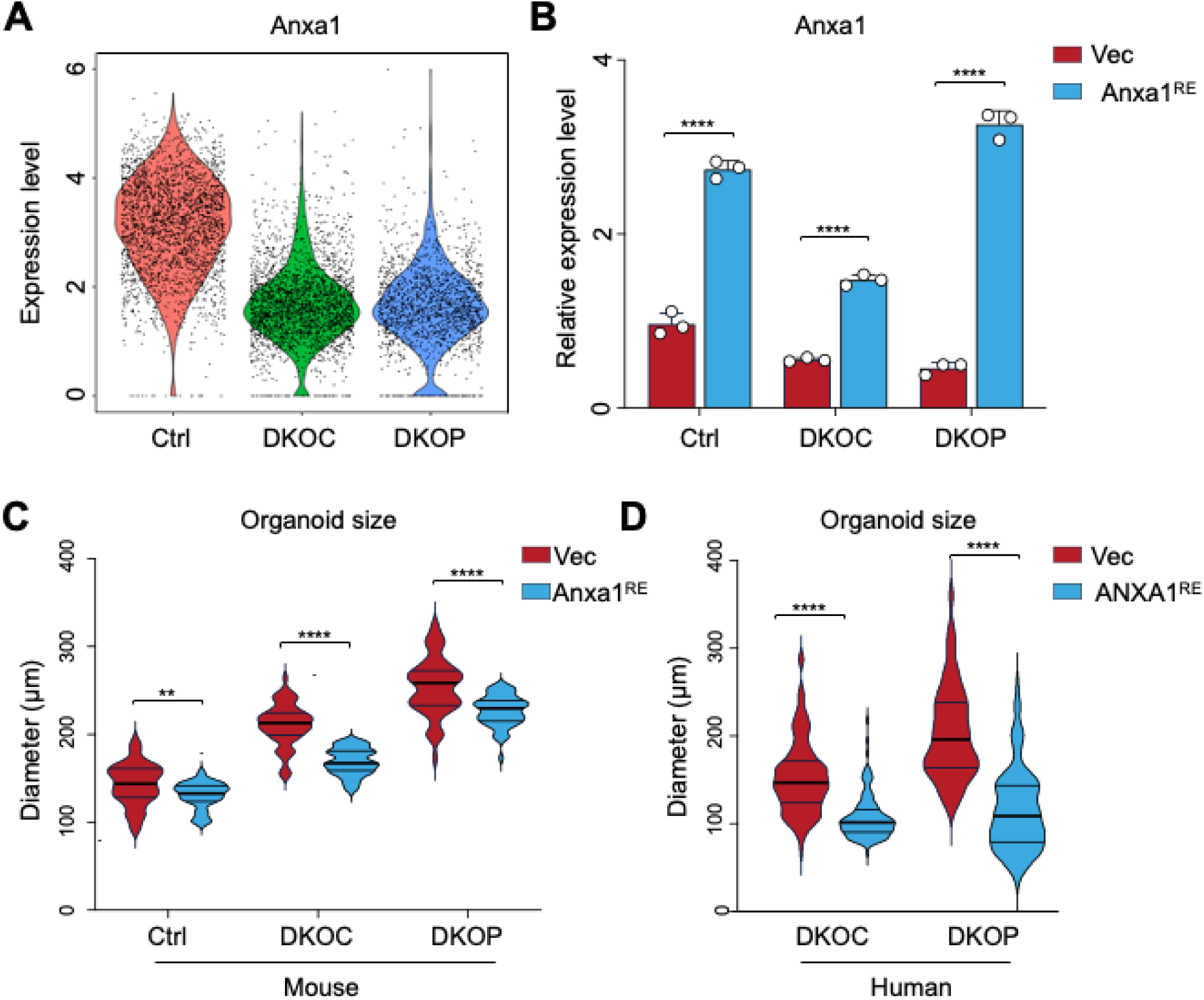
Analysis of ANXA1 expression and growth properties in mouse and human tongue organoids. **(A)** Anxa1 expression in mouse tongue organoids analyzed using scRNA-seq data. **(B)** Relative Anxa1 mRNA expression in Vec and Anxa1^RE^ mouse tongue organoid lines, normalized to the mouse tongue Ctrl Vec group. **(C)** Average size of mouse and human tongue Ctrl, DKOC, and DKOP organoid lines. \*\**P* < 0.01; \*\**P* < 0.0001.

**Figure S12.**
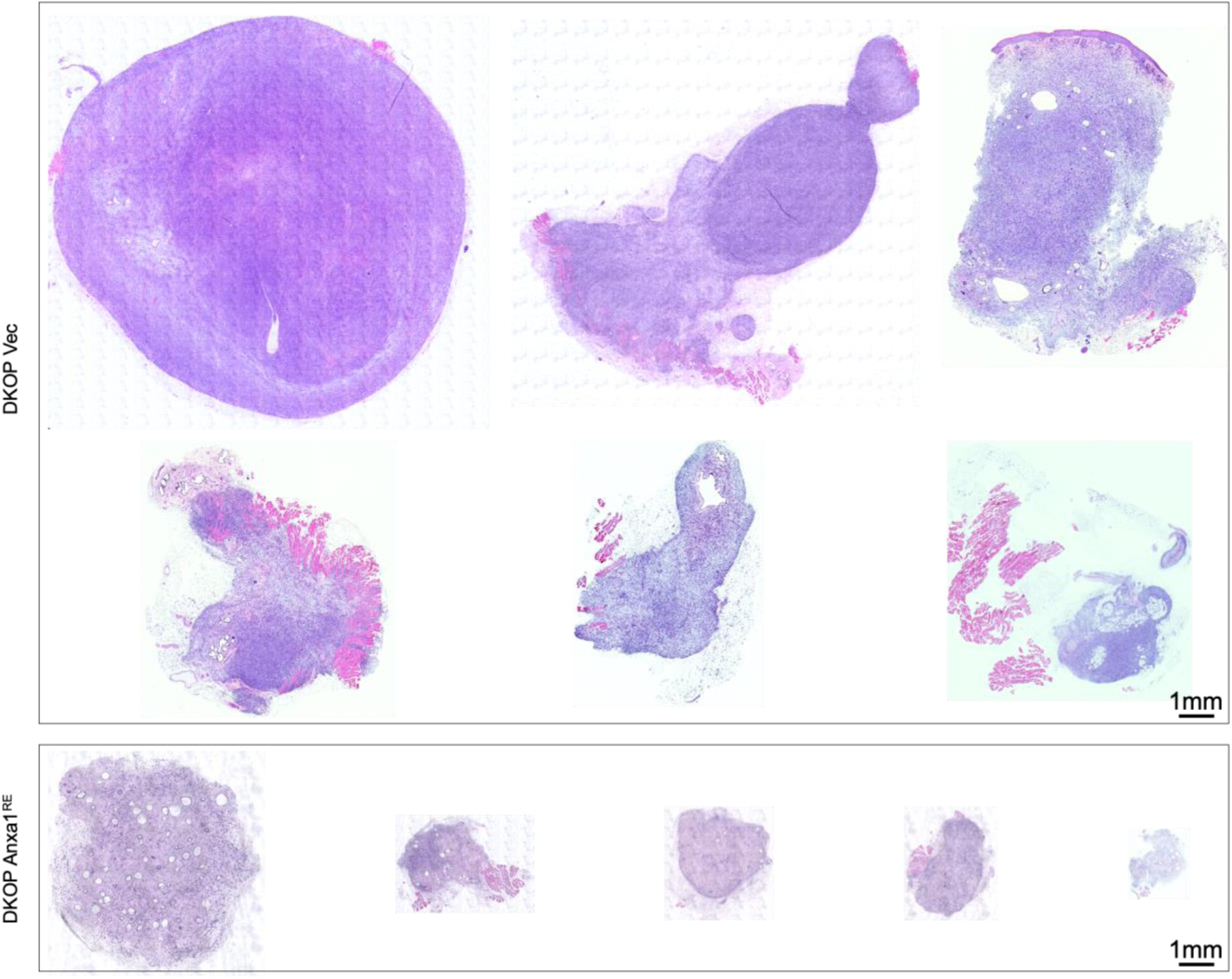
H&E staining showing Anxa1^RE^ in DKOP organoids yields fewer and smaller allografts with reduced atypical features.

**Figure S13.**
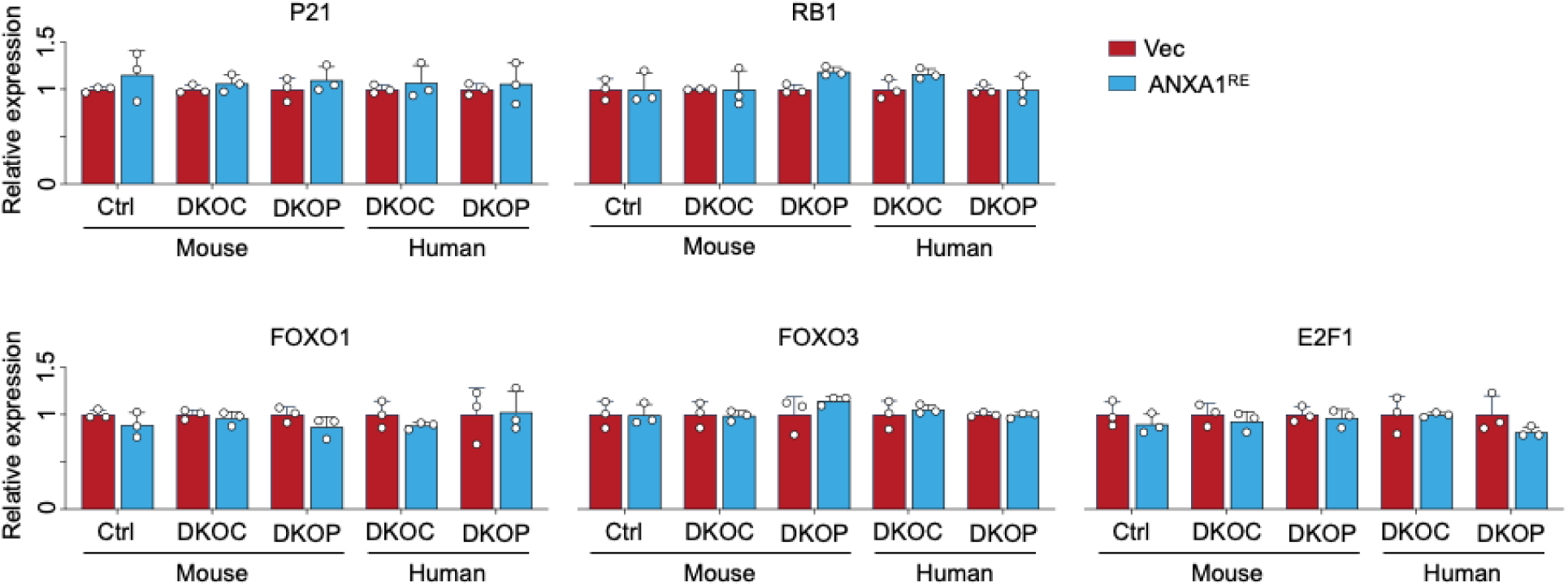
Relative mRNA expression in Vec and Anxa1^RE^ mouse and human tongue organoid lines, normalized to the corresponding Vec group.

**Figure S14.**
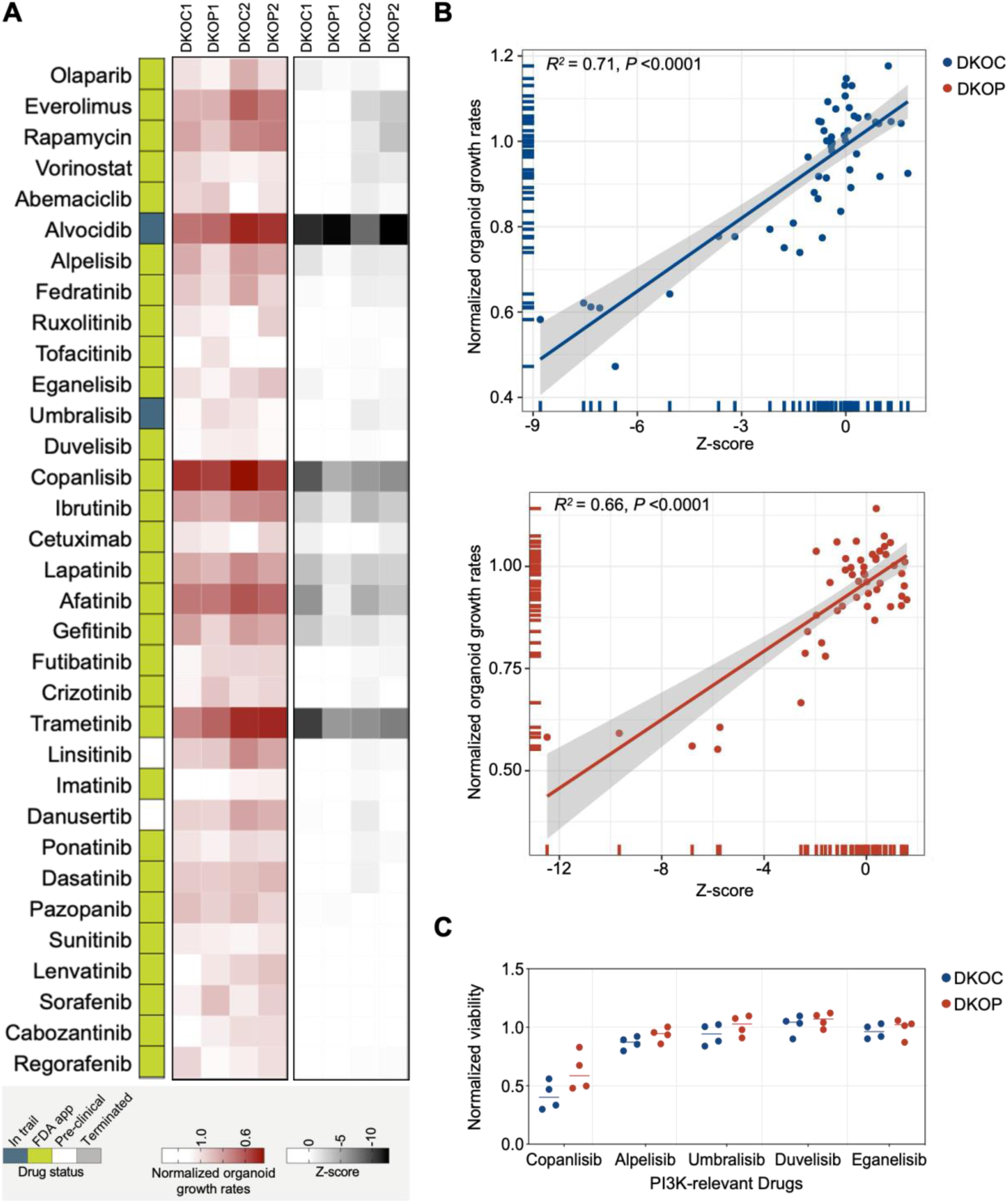
Drug screening and correlation analysis of ATP-based measurements with organoid viability assessed via machine-learning-assisted image analysis. **(A)** Heatmaps of growth rates and ATP assay results (z-scores) for organoids. Growth rates were calculated as the ratio of organoid growth on day 6 (prior to the ATP assay) to day 4 (before the initial drug treatment), normalized to the DMSO vehicle control. Data from each of the two independent biological replicates are shown. **(B)** Scatter plot of linear regression analysis between z-score and organoid growth rates assessed through machine learning-based image analysis. **(C)** Viability of mouse DKOC and DKOP organoids under PI3K-relevant drug treatments, measured by ATP assay.

**Figure S15.**
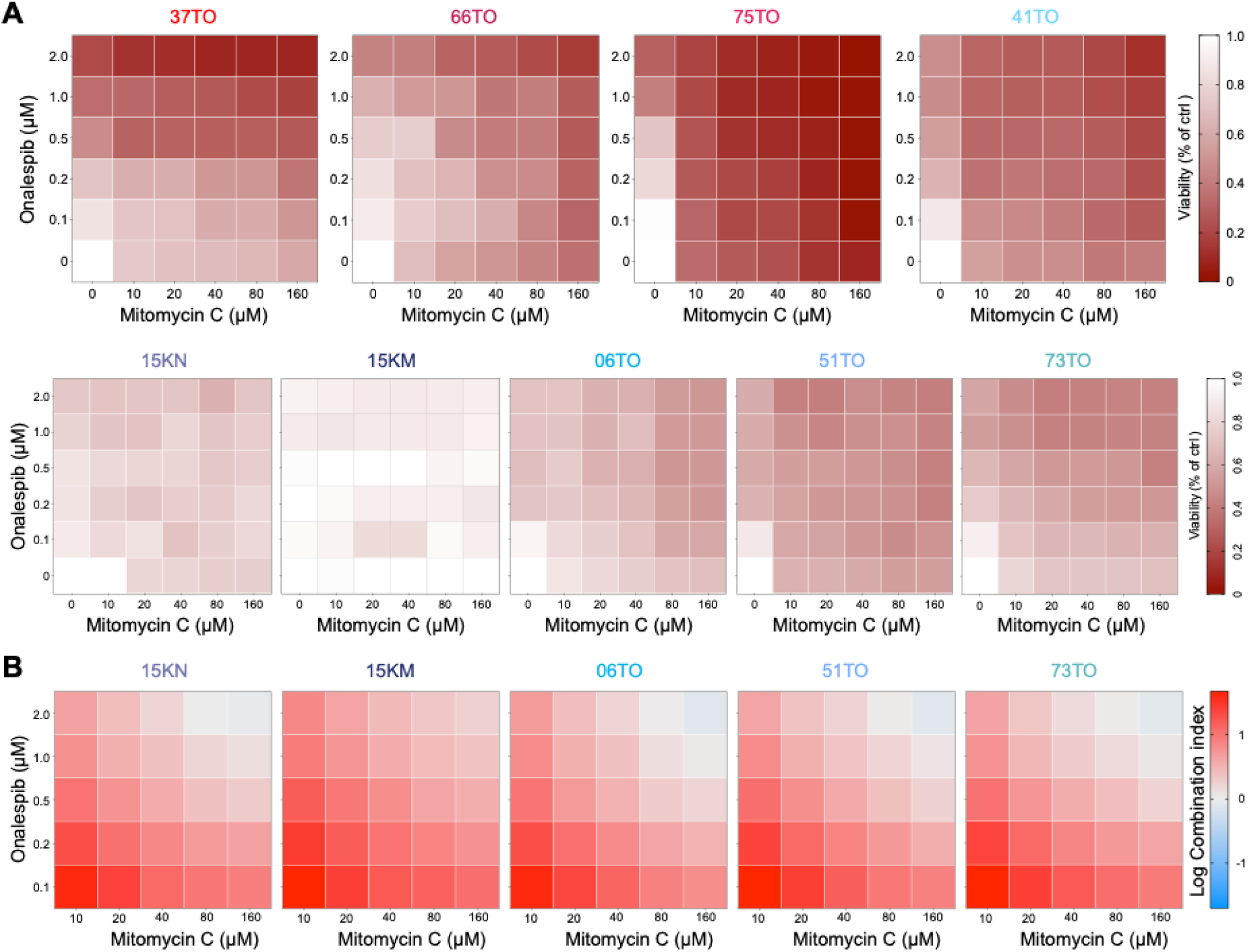
Sensitivity and combination index analysis of Mitomycin C and Onalespib in *PIK3CA* wild-type and mutant PDOs. **(A)** Sensitivity of *PIK3CA-*mutant (37TO, 66TO, and 75TO), *PIK3CA-*amplified (41TO) and wild-type (lower panel) PDOs to the combination of Mitomycin C and Onalespib (n=3). The data is normalized to DMSO control. **(B)** Combination index (CI) of Mitomycin C and Onalespib across wild-type *PIK3CA* PDOs (n=3). Log CI > 0 indicates antagonism; Log CI = 0 indicates additivity; Log CI < 0 indicates synergy.

**Figure S16.**
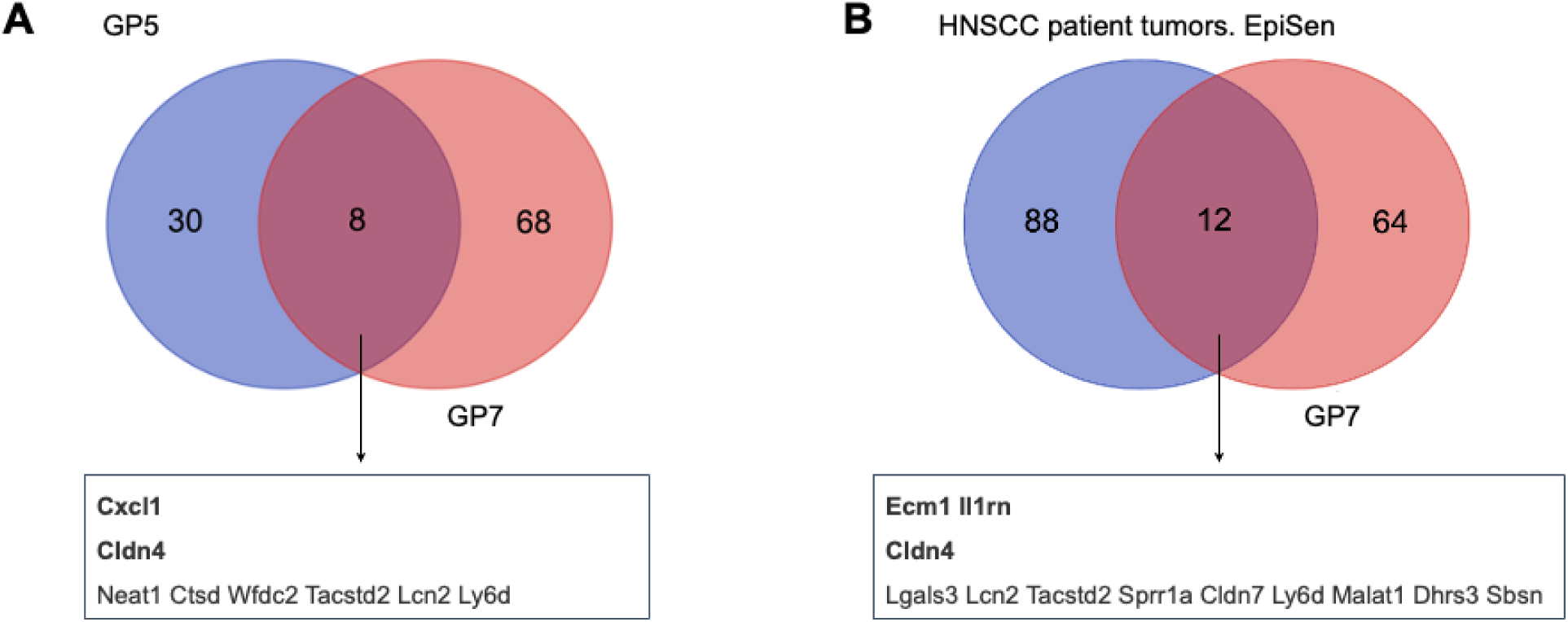
Venn diagram analysis of overlapping genes between mouse organoid GP7 and GP5 (left), and between GP7 and the EpiSen program in HNSCC patient tumors (right).

**Table S1. Clinical information of patient samples**

**Table S2. Key resources table**

**Table S3. Gene list for identified GPs**

